# RBMS1 suppresses colon cancer metastasis through targeted stabilization of its mRNA regulon

**DOI:** 10.1101/2020.01.22.916205

**Authors:** Johnny Yu, Bruce Culbertson, Hosseinali Asgharian, Albertas Navickas, Lisa Fish, John Paolo Olegario, Benjamin Hänisch, Ethan M Weinberg, Rodrigo Dienstmann, Robert S Warren, Hani Goodarzi

## Abstract

Broad dysregulation of gene expression control is a hallmark of cancer progression. Identifying the underlying master regulators that drive pathological gene expression is a key challenge in precision oncology. Here, we have developed a network analytical framework, named PRADA, that identifies oncogenic RNA-binding proteins through the systematic detection of coordinated changes in their target regulons. Application of this approach to data collected from clinical samples, patient-derived xenografts, and cell line models of colon cancer metastasis revealed the RNA-binding protein RBMS1 as a suppressor of colon cancer progression. We observed that silencing RBMS1 results in increased metastatic capacity in xenograft mouse models, and that restoring its expression blunts metastatic liver colonization. We have found that RBMS1 functions as a post-transcriptional regulator of RNA stability by directly binding and stabilizing ~80 target mRNAs. Measurements in more than 180 clinical samples as well as survival analyses in publicly available datasets, have shown that RBMS1 silencing and the subsequent downregulation of its targets are strongly associated with disease progression and poor survival in colon cancer patients. Together, our findings establish a role for RBMS1 as a previously unknown regulator of RNA stability and as a suppressor of colon cancer metastasis with clinical utility for risk stratification of patients.

**Significance:** By applying a new analytical approach to transcriptomic data from clinical samples and models of colon cancer progression, we have uncovered RBMS1 as a suppressor of metastasis and as a post-transcriptional regulator of RNA stability. Notably, RBMS1 silencing and downregulation of its targets are negatively associated with patient survival.

## Main

Metastatic progression in colorectal cancer (CRC) is accompanied by widespread gene expression reprogramming. Cancer cells often co-opt post-transcriptional regulatory mechanisms to achieve pathological expression of gene networks that drive metastasis (1–3). Colorectal cancer is the third most commonly diagnosed cancer (4), therefore understanding the underlying regulatory programs that drive metastatic progression in this disease is a crucial step towards improving patient outcomes. Notably, there are not many predictive computational methods aimed at the discovery of unknown regulatory networks. By relying on annotated post-transcriptional regulatory pathways, e.g. those mediated by microRNAs, existing methods fail to capture previously unknown regulatory interactions. To tackle this problem, we have developed a computational approach called PRADA that identifies post-transcriptional master regulators responsible for aberrant mRNA stability and gene expression in cancer cells. By applying this tool to a large compendium of gene expression profiling data from patient samples, patient-derived xenograft models, and established colon cancer cell lines, we identified a novel regulatory program involved in CRC metastasis. We find that this previously unknown regulatory pathway, which controls mRNA stability and is mediated by the RNA-binding protein RBMS1, is often silenced in highly metastatic CRC tumors. We demonstrate that RBMS1 stabilizes transcripts by binding to the last exons of target mRNAs, and that in highly metastatic CRC cells and patient tumors, RBMS1 downregulation is associated with poor clinical outcome. We identify mRNA targets of RBMS1 that are functionally relevant to metastasis and reveal that RBMS1 silencing can be accomplished through epigenetic dysregulations. This study not only describes a disease-relevant post-transcriptional regulatory mechanism that governs the stability of a sizeable regulon, but also demonstrates the value of bottom-up discovery strategies like PRADA that do not solely rely on prior knowledge of annotated regulatory programs.

## Results

### PRADA identifies RBMS1 as a novel regulator of CRC progression

Metastasis is a complex multistep process and requires modulation of many cellular pathways and processes. As such, increased metastatic capacity often involves broad reprogramming of the gene expression landscape in cancer cells. Thus, a mechanistic dissection of cancer progression relies on the systematic identification of the underlying regulatory programs that drive pathologic cellular states. To accomplish this, we developed PRADA (Prioritization of Regulatory Pathways based on Analysis of RNA Dynamic Alterations) that uses regulatory network predictions to identify the key RNA-binding proteins (RBPs) that may act as master regulators of mRNA stability and gene expression. PRADA solves a regression problem to predict changes in gene expression as a function of the expression of RBPs and their predicted targets across the transcriptome. In this customized regression analysis, the coefficient assigned to each RBP reflects its strength as a regulator of gene expression and the direction of its effect (i.e. activator or repressor). Given our limited knowledge of post-transcriptional regulatory pathways and their role in human disease, PRADA provides an opportunity for the discovery of previously unknown disease pathways. To assess the contribution of post-transcriptional regulatory programs to colon cancer metastasis, we took advantage of a publicly available set of gene expression profiles from established colon cancer cell lines (GSE59857). We first categorized these cell lines as poorly metastatic or highly metastatic based on their liver colonization capacity in xenograft mouse models (Supplementary Fig. S1A; (5)). We then performed differential gene expression analysis to compare gene expression changes across the transcriptomes of the two groups. Finally, we applied PRADA to this dataset to find RNA-binding proteins whose differential activity is most informative of metastasis-associated gene expression changes. PRADA identified the protein RBMS1 as the RNA-binding protein with the strongest regulatory potential in this dataset (Fig. 1A). The size and the direction of the regulatory coefficient assigned to RBMS1 implies that RBMS1 levels are strongly informative of changes in the expression of transcripts with matches to RBMS1 binding sites. To confirm this, we performed motif enrichment analysis using FIRE, which uses mutual information to capture the association between the presence and absence of a given binding site and genome-wide transcriptomic measurements (6). As shown in Fig. 1B, the RBMS1 consensus binding site is strongly informative of gene expression differences between poorly and highly metastatic cells, showing a highly significant enrichment among the transcripts that have decreased expression in highly metastatic cells. To ensure that this result is not sensitive to the choice of a specific RBMS1 binding motif, we tested two additional sequence models: (i) an independently derived representation of RBMS1 binding preferences (7), and (ii) predictions based on DeepBind models (8). Each of these three predicted RBMS1 binding motifs gave largely identical results, and we observed a similar decrease in the expression of the putative RBMS1 regulon derived from each motif (Supplementary Fig. 1B). For consistency, we have used the DeepBind RBMS1 model in our subsequent analyses as it is the state-of-the-art approach for predicting protein-nucleic acid interactions. Importantly, consistent with the results reported by PRADA and the lower expression of its putative regulon, the expression of RBMS1 was also significantly lower in highly metastatic cells (Fig. 1C). Moreover, consistent with RBMS1 acting as a post-transcriptional regulator of these genes, we observed a significant correlation between the expression of RBMS1 and the average expression of its regulon in multiple independent datasets (Fig. 1D).

**Figure 1.**
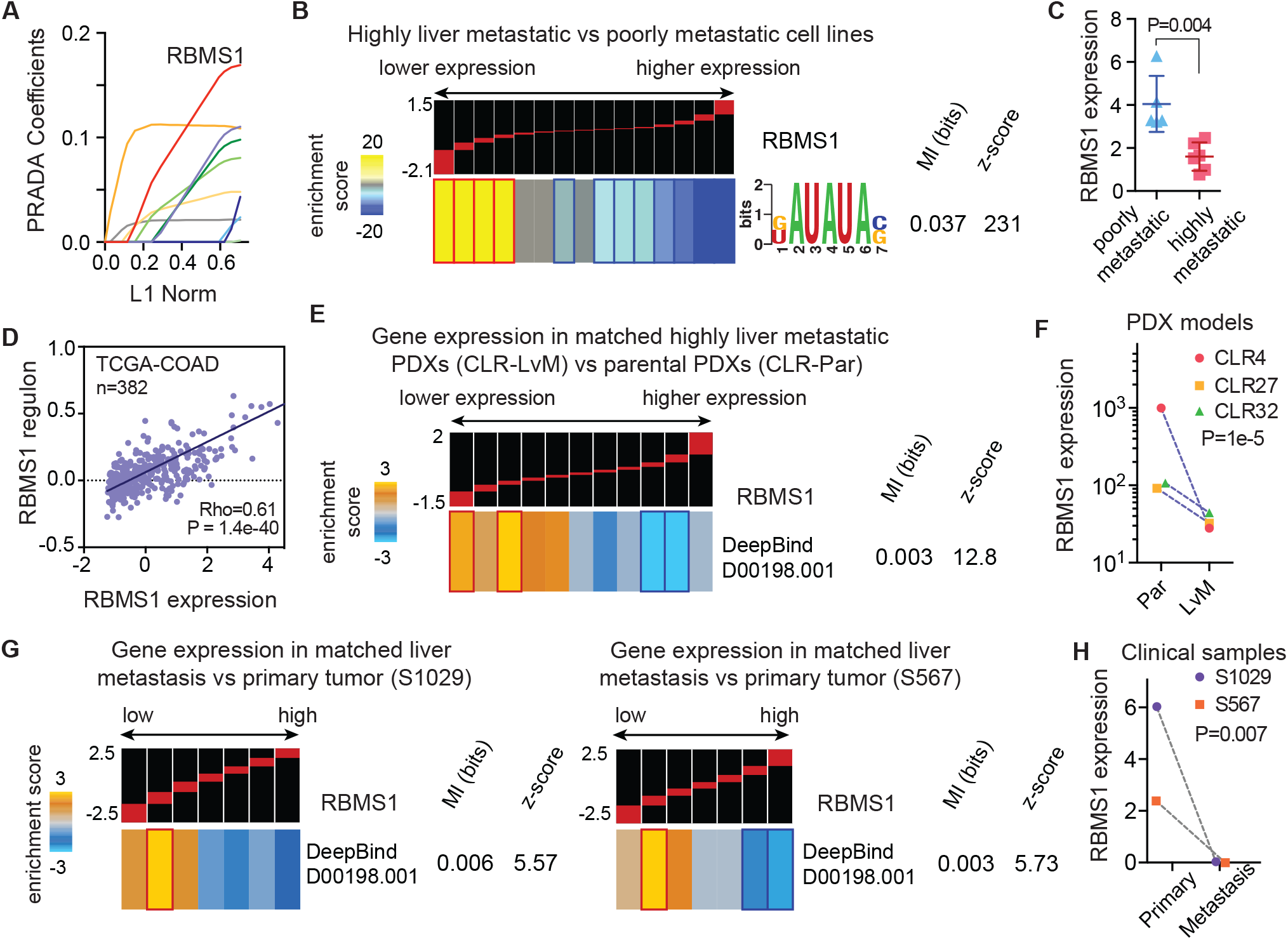
RBMS1 silencing in metastatic cells is associated with lower expression of RBMS1 targets. **(A)** Regression coefficients set by PRADA as a function of the *l1*-norm of the entire coefficient vector. Each line is associated with an RNA-binding protein and the magnitude of its coefficient is a measure of its strength as a putative regulator of gene expression. Here, the first ten non-zero coefficients are shown as a function of *l1*-norm (i.e. sum of the magnitude of all coefficients). The red line represents the coefficient set for the RNA-binding protein RBMS1 at each iteration. **(B)** Analysis of RBMS1 recognition sites across gene expression changes between poorly and highly metastatic colon cancer lines. In this analysis, transcripts are first ordered based on their log-fold changes from left (lower expression in metastatic cells) to right (higher expression) and then partitioned into equally populated bins (~1000 transcripts per expression bin). The red bars on the black background show the range of values in each bin (with the minimum and maximum values, i.e. −1.5 and 2, presented on the left). As shown here, genes that are expressed at a lower level in highly metastatic cells were significantly enriched for the RBMS1 binding motif (KAUAUAS)(10). In this heatmap, gold represents overrepresentation of putative RBMS1 targets while blue indicates underrepresentation. Enrichment and depletions that are statistically significant (based on hypergeometric distribution) are marked with red and dark blue borders, respectively. Also included are the logo representation of the RBMS1 binding motif, its mutual information (MI) value and the associated z-score (for details of this analysis see (11)). **(C)** RBMS1 expression in colon cancer lines grouped based on their metastatic capacity (Supplementary Fig. 1A). *P* calculated based on a two-tailed Mann-Whitney *U*-test. **(D)** Linear regression analysis of RBMS1 expression versus that of its putative target signature in TCGA-COAD dataset (cbioportal; N = 382). Shown are the Spearman correlation coefficient and the associated *p*-value. **(E)** Enrichment and depletion patterns of the RBMS1 regulon in PDX models of CRC liver metastasis. For this analysis, log-fold changes between parental (CLR-Par) and liver metastatic (CLR-LvM) were averaged across three independent PDX models, CLR4, CLR27, and CLR32. The distribution of RBMS1 targets, as determined by DeepBind (8), was then assessed using mutual information and its associated *z*-score. The enrichment and depletion patterns were visualized as described in (**B**). **(F)** Relative RBMS1 levels in matched poorly metastatic (Par) and highly liver metastatic (LvM) derivatives for three PDX models (CLR4, CLR32, and CLR27). *P* was calculated using DESeq2 (12). **(G-H)** Expression of RBMS1 and RBMS1 targets in matched primary tumor and liver metastases from two patients (S1029 and S567). The results are presented as described in (**B**). The *P* value for RBMS1 silencing was calculated using DESeq2.

In order to further assess the association between RBMS1 and CRC metastasis in more directly disease-relevant models, we took advantage of patient-derived xenograft (PDX) models of colon cancer metastasis to liver. We used RNA-seq data from three parental CRC PDX models (CLR4, CLR27, and CLR32) and their highly liver metastatic derivatives (CLR-LvMs). The highly metastatic CLR-LvM models were derived from the parental PDXs by repeated splenic delivery and growth of cells in the liver of immunocompromised mice (NOD scid gamma), recapitulating the major site of CRC metastasis in humans (9). We observed that the increase in the metastatic capacity of the CLR-LvM PDX models compared to their parental PDXs was accompanied by a significant reduction in the expression of the RBMS1 regulon in the highly metastatic CLR-LvM models (Fig. 1E). More importantly, all three models also showed a significant and concomitant decrease in the expression of RBMS1 (Fig. 1F).

In addition to these cell line and PDX models of CRC liver metastasis, we also performed RNA-seq on matched primary tumors and liver metastases biopsied and frozen from two patients. As shown in Fig. 1G, in both cases, the RBMS1 regulon was enriched among transcripts that were downregulated in the metastases relative to their primary tumors. Interestingly, while the gene expression changes between the primary tumors and metastases were not generally correlated (R = 0.02, P = 0.1), the RBMS1 regulon was independently downregulated in both metastatic tumors (Supplementary Fig. 1C). Consistently, RBMS1 was also strongly silenced in both metastases (Fig. 1H). Collectively, these findings in cell lines, PDXs, and clinical samples implicate RBMS1 silencing in both the downregulation of its putative regulon and in CRC metastatic progression.

### RBMS1 acts as a post-transcriptional regulator of RNA stability

The correlated expression of RBMS1 and its putative regulon defined from its binding site preferences implies that RBMS1 modulates gene expression. To further assess the potential role of RBMS1 as a post-transcriptional regulator, we performed RNA sequencing of RBMS1 knockdown and control SW480 colon cancer cells. We chose the SW480 cell line for this experiment because: (i) RBMS1 expression in SW480 is among the highest in colon cancer lines tested by us and others, and (ii) it is an established xenograft model of liver metastasis and is considered to be poorly metastatic (relative to other CRC lines listed in Supplementary Fig. 1A). We used two independent shRNAs to silence RBMS1 in SW480 cells and confirmed its knockdown with both qPCR and Western blotting (Supplementary Fig. 2A). As shown in Fig. 2A, a ~4-fold reduction in RBMS1 expression resulted in a significant decrease in the expression of its regulon. Since we are focused on the role of RBMS1 as an RNA-binding protein, we reasoned that its effect on gene expression is likely through regulation of the stability of its target RNAs. To test this hypothesis, we took advantage of REMBRANDTS, a computational framework we have developed to estimate RNA stability from RNA-seq data (13). We have previously established that REMBRANDTS accurately recapitulates experimental RNA stability measurements (13). Application of this method to RNA-seq data from RBMS1 knockdown and control cells found a significant enrichment of RBMS1 targets among transcripts destabilized upon RBMS1 silencing (Fig. 2B). REMBRANDTS relies on the comparison of exonic and intronic reads to measure changes in RNA stability. As shown in Supplementary Fig. 2B, RBMS1 silencing results in lower expression of its putative regulon as measured by exonic reads; however, intronic reads, which reflect changes in pre-mRNA levels (and transcription rates) do not significantly change in response to RBMS1 depletion. To further strengthen our claim that RBMS1 functions as a post-transcriptional regulator of RNA stability rather than as a transcriptional activator, we also performed whole-genome RNA stability measurements in control and RBMS1 knockdown cells. For this, we used α-amanitin-mediated inhibition of RNA-polymerase II followed by RNA sequencing at two time points (0-hr and 9-hr). Consistent with our analyses with REMBRANDTS, we observed a similar reduction in the stability of the putative RBMS1 regulon upon RBMS1 silencing (Fig. 2C). We used RNA-seq data from the three matched PDX models (CLR panel) to compare RNA stability in highly liver metastatic models to that of their parental PDXs (using REMBRANDTS). As shown in Fig. 2D, lower RBMS1 expression in the LvM models (Fig. 1F) accompanies a reduction in the stability of its target regulon in all three independently derived models. Together, these findings establish RBMS1 as a post-transcriptional regulator of RNA stability with functional implications for CRC progression.

**Figure 2.**
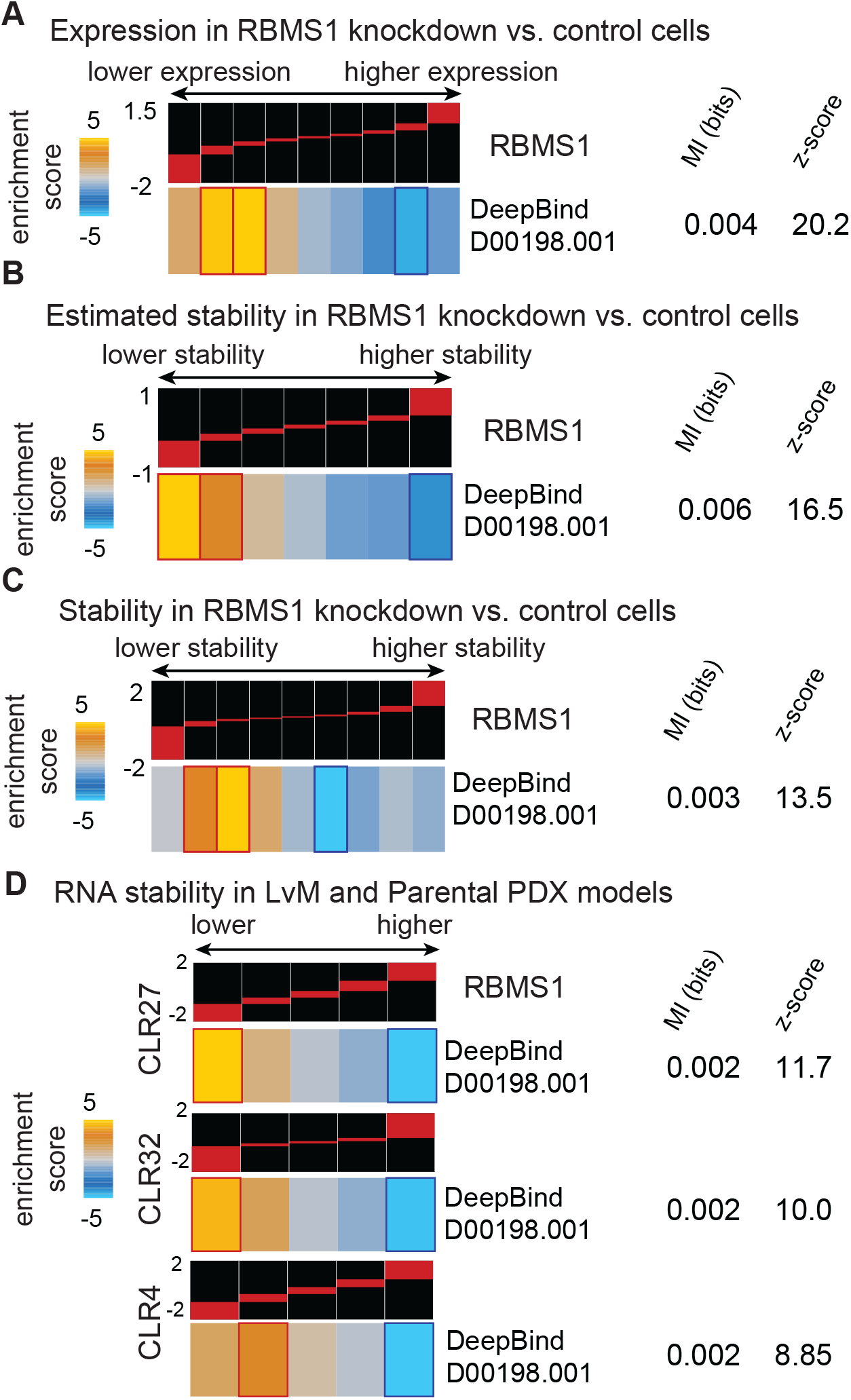
RBMS1 post-transcriptionally regulates the stability and expression of its targets. **(A)** Enrichment and depletion patterns of the RBMS1 regulon in RBMS1 knockdown cells relative to control (~4-fold knockdown). **(B)** We used our computational tool, called REMBRANDTS, to estimate changes in RNA stability upon RBMS1 silencing. These differential stability estimates were then used to assess the enrichment patterns of the RBMS1 targets across the changes in RNA decay. **(C)** Experimental RNA stability changes were measured using α-amanitin treatment as previously described (14). The enrichment and depletion patterns of the RBMS1 regulon was then assessed among the transcripts destabilized or stabilized upon RBMS1 knockdown. **(D)** We used REMBRANDTS to measure changes in RNA stability between poorly and highly metastatic PDX models from three independent PDX models (CLR27, CLR32, and CLR4). As shown here, consistent with the silencing of RBMS1 in LvM PDX models (Fig. 1F) and the down-regulation of its regulon (Fig. 1E), the RBMS1 regulon is destabilized in these three independent models of CRC metastasis.

### RBMS1 is a suppressor of EMT and metastatic liver colonization in colon cancer cells

In order to carry out a functional study of RBMS1 and its downstream regulon in CRC metastasis, we picked an additional cell line model to complement the SW480 line. We chose the LS174T line because RBMS1 is almost completely silenced in this line as measured by qPCR and Western blotting (Fig. S3A-B). Also, LS174T is an established xenograft model and is considered a highly liver metastatic cell line, ~100x more metastatic than the SW480 line as measured by liver colonization assays (15). We first performed RNA sequencing of both SW480 and LS174T lines and consistent with RBMS1 silencing in the LS174T cells, we observed a significant reduction in the expression of the RBMS1 putative regulon in this line (Supplementary Fig. 3C).

To establish a causal link between RBMS1 silencing and higher metastatic capacity, we performed liver colonization assays by splenically injecting RBMS1 knockdown and control SW480 cells and measuring metastatic burden in the livers of mice over time. While RBMS1 silencing did not have a strong effect on *in vitro* cell proliferation (Supplementary Fig. 3D), we observed a significant increase in liver colonization upon silencing RBMS1, based on *in vivo* bioluminescence measurements and gross liver mass at the experimental endpoint (Fig. 3A; Supplementary Fig. 3E). To control for possible off-target effects of the shRNAs, we repeated the experiment with an additional RBMS1-targeting hairpin and observed a similarly significant increase in metastatic liver colonization (Supplementary Fig. 3F). We also performed the converse gain-of-function experiment by expressing RBMS1 in the LS174T line, where RBMS1 is endogenously silenced. Consistent with our earlier findings, we observed that expressing RBMS1 in LS174T cells results in a marked reduction in their liver colonization capacity (Fig. 3B; Supplementary Fig. 3G). It should be noted that as expected, expressing shRNAs targeting RBMS1 in highly metastatic cells in which RBMS1 is already silenced, namely LS174T, HCT116, and Colo320, did not have an impact on their liver colonization capacity in xenograft mouse assays, which further highlights the on-target effect of the assayed shRNAs (Supplementary Fig. 3H). In contrast, knockdown of RBMS1 in WiDr cells, in which RBMS1 expression is intact, resulted in a significant increase in liver colonization in xenograft mouse models without an overt effect on *in vitro* cell proliferation (Supplementary Fig. 3I-J).

**Figure 3.**
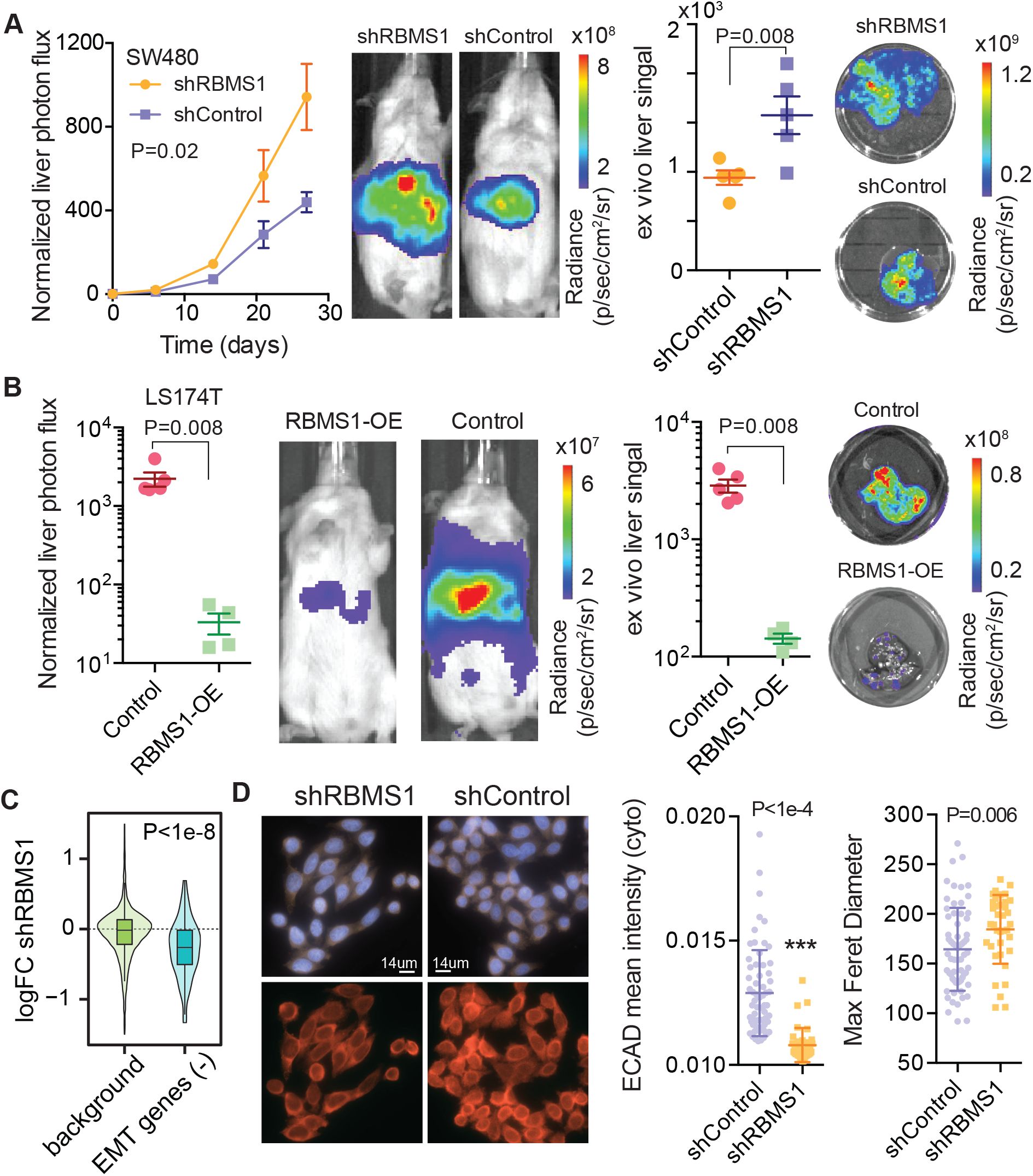
RBMS1 is a suppressor of epithelial-mesenchymal transition (EMT) and metastatic liver colonization. **(A)** Bioluminescence imaging plot of liver colonization by RBMS1 knockdown or control cells; N = 5 in each cohort. Two-way ANOVA was used for statistical testing. *Ex vivo* liver signal was also measured and compared using a one-tailed Mann-Whitney *U*-test. Also shown are representative (median signal) mice and livers. **(B)** Splenic injection of LS174T highly metastatic colon cancer cells overexpressing RBMS1 and those expressing mCherry as a control. Day 21 signal (normalized to day 0) was plotted and compared for both *in vivo* and *ex vivo* signal (N = 4-5; Mann-Whitney *U*-test). **(C)** Downregulation of EMT(−) signature genes upon RBMS1 knockdown in SW480 cells. EMT(−) genes were compared to the rest of the transcriptome using a Mann-Whitney *U*-test. **(D)** Immunofluorescence staining for E-Cadherin (red) in RBMS1 knockdown and control cells (SW480 background). Note the lower expression of E-Cadherin and the more spindle-like cellular morphology in RBMS1 knockdown (top panels show DAPI signal). ECAD intensity and maximum Feret Diameter (a measure of length of the cell) for cells in control and knockdown samples (N = 64 and 37, respectively). Two-tailed Mann-Whitney *U*-test was used to compare measurements.

The metastatic cascade is a complex multistep process, involving cellular processes that not only impact cancer cell growth and survival, but also cell migration and invasion. As mentioned earlier, we did not observe a significant change in *in vitro* cell proliferation rates upon RBMS1 knockdown. Additionally, trans-well invasion assays of RBMS1 knockdown and control cells did not detect a significant role for RBMS1 in cancer cell invasion (Supplementary Fig. 3K). Therefore, to gain a better understanding of the key step(s) in the metastatic cascade that are regulated by RBMS1, we performed a systematic analysis of known and predicted gene-sets to identify those that may be modulated upon RBMS1 silencing. We performed this analysis using iPAGE, an information-theoretic framework we have developed for this type of analysis (16). Downregulation of genes associated with negative regulation of epithelial-mesenchymal transition (160-gene EMT(−) signature (17)) was the most significant pathway identified in RBMS1 knockdown cells (Fig. 3C; Supplementary Fig. 3L). For example, the canonical EMT marker E-Cadherin is significantly downregulated by ~2-fold in RBMS1 knockdown cells based on RNA-seq data. We confirmed this by performing immunofluorescence staining for E-Cadherin in RBMS1 knockdown and control cells. We observed both a reduction in E-cadherin expression in RBMS1 knockdown cells, and also noted the appearance of spindle-like cell morphology that is associated with EMT (Fig. 3D). Moreover, the EMT(−) signature genes were also expressed at relatively lower levels in the LS174T cell line, where RBMS1 is endogenously silenced (Supplementary Fig. 3L). Finally, analysis of the TCGA-COAD dataset revealed significantly lower E-cadherin expression in tumor samples with low RBMS1 expression (~20% reduction, *P*=0.006) and a significant general correlation between RBMS1 and E-Cadherin expression (Rho=0.1, P=0.03), which is indicative of the clinical relevance of this regulatory axis in CRC.

### RBMS1 CLIP-seq reveals 3’UTR binding of target mRNAs

Our results indicate that RBMS1 acts as a post-transcriptional regulator of RNA stability and its silencing results in the downregulation of its putative regulon. As mentioned above, these analyses were performed using *in silico* predicted binding targets of RBMS1. In order to create a transcriptome-wide snapshot of RBMS1 binding sites in colon cancer cells at nucleotide resolution, we performed UV crosslinking immunoprecipitation followed by sequencing (CLIP-seq). We carried out irCLIP (18) for endogenous RBMS1 in SW480 cells, and identified hundreds of high-confidence RBMS1 binding sites across the transcriptome (Fig. 4A). Importantly, we observed that 90% of the CLIP-identified RBMS1 regulon overlapped with our computationally-derived putative RBMS1 regulon. Consistently, these RBMS1 targets (defined as RNAs bound by RBMS1 in our irCLIP data) follow the same expression patterns as the computationally-derived RBMS1 regulon described above, exhibiting both decreased stability and expression upon RBMS1 silencing (Fig. 4B; Supplementary Fig. 4A).

**Figure 4.**
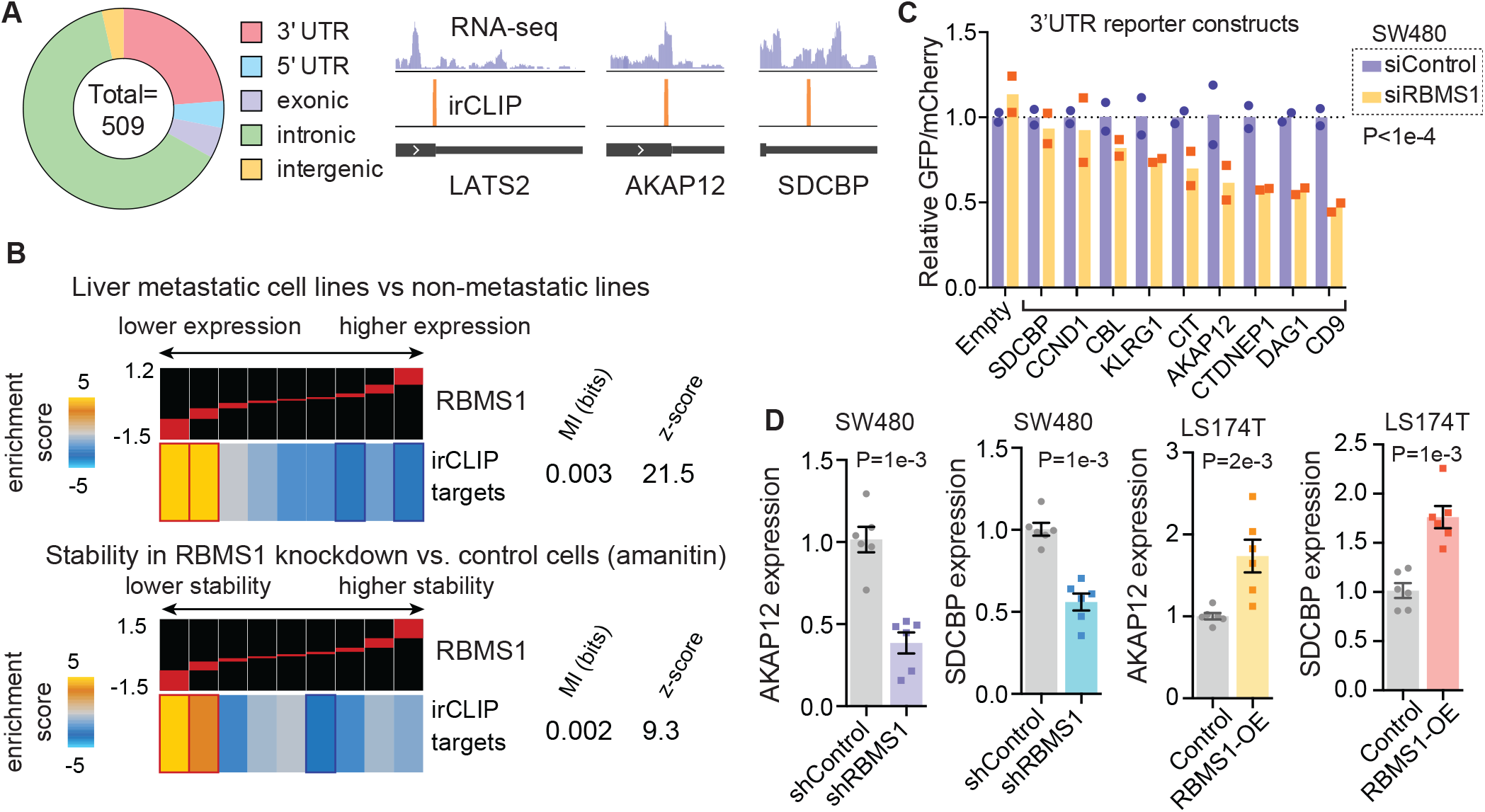
RBMS1 irCLIP identifies direct RBMS1 targets in colon cancer cells. **(A)** 509 RBMS1 binding sites were found using irCLIP, with a significant enrichment of binding to the last exon/3’UTR (relative to the total length of genomic features). Last exons from LATS2, AKAP12, and SDCBP are shown as examples of RBMS1 binding patterns. **(B)** Enrichment of the RBMS1-bound mRNAs among those that are downregulated in highly metastatic cells (top) and those destabilized upon RBMS1 silencing (bottom). **(C)** qRT-PCR was used to measure changes in GFP mRNA levels upon cloning RBMS1 binding sites of the listed genes downstream of the GFP ORF. mCherry was expressed from the same bidirectional promoter as GFP, and mCherry levels were used to normalize GFP measurements. A one-sample Wilcoxon signed rank test was used to assess whether the ratios in siRBMS1 samples were significantly below 1.0. **(D)** Changes in the expression of AKAP12 and SDCBP upon RBMS1 knockdown in SW480 cells and RBMS1 over-expression in LS174T cells.

In our analysis of RBMS1 binding sites, we noted a strong enrichment of RBMS1 binding to the last exon (on or close to 3’UTRs; Fig. 4A). This is consistent with RBMS1 acting as a post-transcriptional regulator of RNA stability, and suggested that RBMS1 3’UTR binding could result in increased RNA stability. To assess this possibility, we built a bi-directional CMV promoter that drives the expression of both GFP and mCherry. We then cloned nine CLIP-seq-derived RBMS1 binding sites, plus ~150 nucleotides flanking the binding sites, downstream of GFP, and asked whether there was a reduction in GFP mRNA relative to mCherry mRNA upon RBMS1 depletion. As shown in Fig. 4C, there was a reduction in GFP reporter transcript levels in almost all instances. To further ensure that this effect was RBMS1-dependent, we also tested the two reporters with the strongest reduction in transcript levels (DAG1 and CD9) in LS174T cells, where RBMS1 is endogenously silenced. In this instance, as expected, there was no response in reporter mRNA levels upon transfection of RBMS1-targeting siRNAs (Supplementary Fig. 4B). Together, these results indicate that direct binding of RBMS1 to mRNA 3’UTRs results in increased mRNA stability.

### AKAP12 and SDCBP are functional downstream targets of RBMS1

In order to identify genes that are regulated by RBMS1 and act as suppressors of metastasis in CRC, we used an integrated analytical approach, which relies on mining relevant datasets to prioritize target genes based on their direct interaction with RBMS1 (3’UTR binding), their RBMS1-dependent magnitude of change in expression, the robustness of their response to RBMS1 silencing, as well as their lower expression in metastatic clinical samples (Supplementary Fig. 4C). Using this approach, we prioritized two target genes, AKAP12 and SDCBP, that satisfy these criteria: (i) destabilized and downregulated in highly metastatic CRC cells and PDXs, (ii) destabilized and downregulated in RBMS1 knockdown cell lines, (iii) directly bound by RBMS1 based on our irCLIP data (Fig. 4A), and (iv) downregulated in liver metastases relative to primary colon cancers in a publicly available dataset (19). Consistently, silencing RBMS1 in SW480 cells resulted in the downregulation and destabilization of AKAP12 and SDCBP mRNAs (Fig. 4D; Supplementary Fig. 4D). Conversely, overexpression of RBMS1 in LS174T cells resulted in upregulation and stabilization of these targets (Fig. 4D; Supplementary Fig. 4D). These observations establish AKAP12 and SDCBP as bona fide direct targets of RBMS1 that are post-transcriptionally regulated by this RBP.

To test the possible role of AKAP12 and SDCBP in driving CRC metastatic progression, we silenced them individually in SW480 cells using CRISPR interference (CRISPRi) and performed liver colonization assays. As shown in Supplementary Fig. 5A, silencing these genes did not have a significant impact on *in vitro* proliferation of SW480 cells. However, reduced expression of these genes resulted in increased metastatic liver colonization in mice (Fig. 5A). To confirm that AKAP12, which, of the two targets tested, elicited a stronger phenotype when silenced, indeed functions downstream of RBMS1, we performed an *in vivo* epistasis experiment by generating cells with knockdown of both AKAP12 and RBMS1 in SW480 cells. As shown in Supplementary Fig. 5B, unlike in Fig. 5A, further silencing of AKAP12 failed to increase metastatic liver colonization in RBMS1 knockdown cells, in which AKAP12 is already downregulated. Moreover, consistent with AKAP12 acting downstream of RBMS1, gene expression profiling of AKAP12 knockdown and control cells revealed a significant induction of an EMT signature (Fig. 5B). Taken together, our findings demonstrate that this RBMS1-AKAP12 regulatory axis acts as a suppressor of EMT and liver metastasis in models of CRC progression.

**Figure 5.**
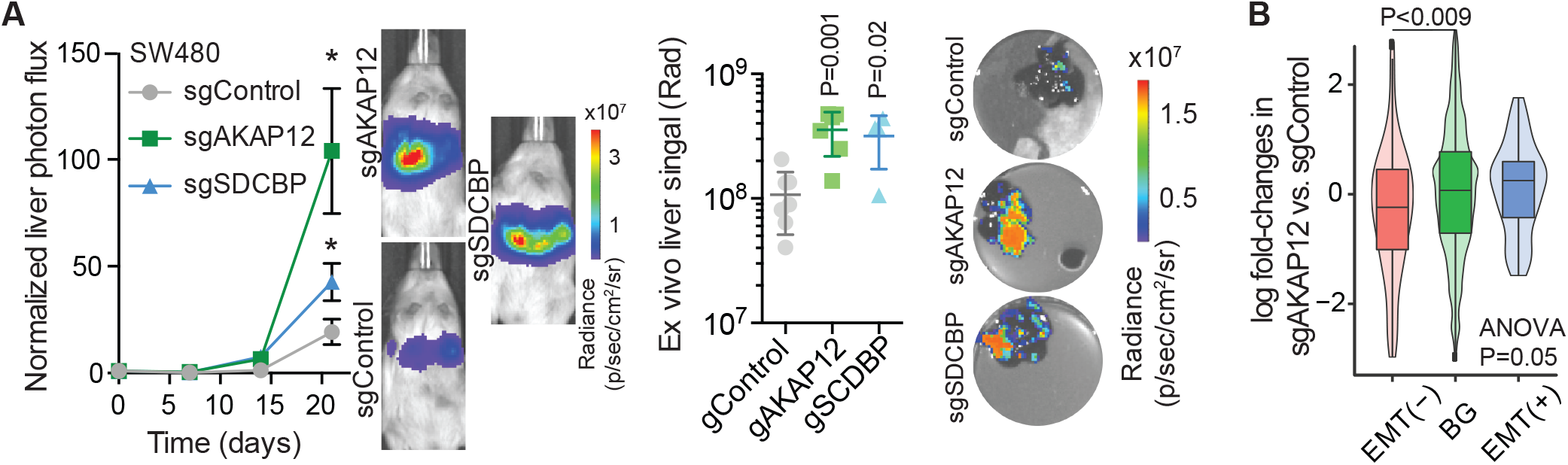
AKAP12 and SDCBP act downstream of RBMS1 to suppress metastasis. **(A)** *In vivo* liver colonization assays were used to measure the impact of CRISPRi-mediated silencing of the RBMS1 targets AKAP12 and SDCBP on liver metastasis (N = 6). Also shown are representative mice from each cohort. Two-way ANOVA was used to compare cohorts to control (P = 0.01 and 0.02 for sgAKAP12 and sgSDCBP respectively). Livers were also extracted and their tumor burden was measured *ex vivo.* Mann Whitney U test was used to compare measurements. **(B)** The expression of EMT(−) and EMT(+) signature genes relative to background (BG) in AKAP12 knockdown (CRISPRi) and control cells (measurements using 3’-end RNA-seq). Shown are the ANOVA *p-*value and a Mann-Whitney comparison between EMT(−) and background genes.

### RBMS1 silencing and the downregulation of its regulon is associated with CRC progression

To further assess the clinical relevance of this previously unknown regulatory pathway, we performed a variety of measurements in clinical samples as well as analysis of clinical data to evaluate RBMS1 activity in CRC metastasis. First, we performed qRT-PCR for RBMS1 in two independent clinical cohorts, one cohort stratifying patients based on their tumor stage (n=96), and another comparing samples from normal mucosa, primary CRC tumors, and liver metastases (n=91). As shown in Fig. 6A and 6B, we observed a significant reduction in RBMS1 expression as the disease progresses, with stage IV clinical samples showing the lowest RBMS1 expression levels. We also carried out survival analysis in a large clinical cohort with publicly available expression data and clinical outcomes (20). We observed a significant association between RBMS1 silencing and reduced relapse-free survival as well as overall survival in colon cancer patients (Supplementary Fig. 6A). A multivariate Cox proportional-hazards model revealed that this association with RBMS1 silencing remains statistically significant even when controlled for other known clinical factors that may contribute to patient survival (Fig. 6C). This observation emphasizes the relevance of RBMS1 silencing as an effective read-out for risk stratification of patients based on samples collected from their primary tumors.

**Figure 6.**
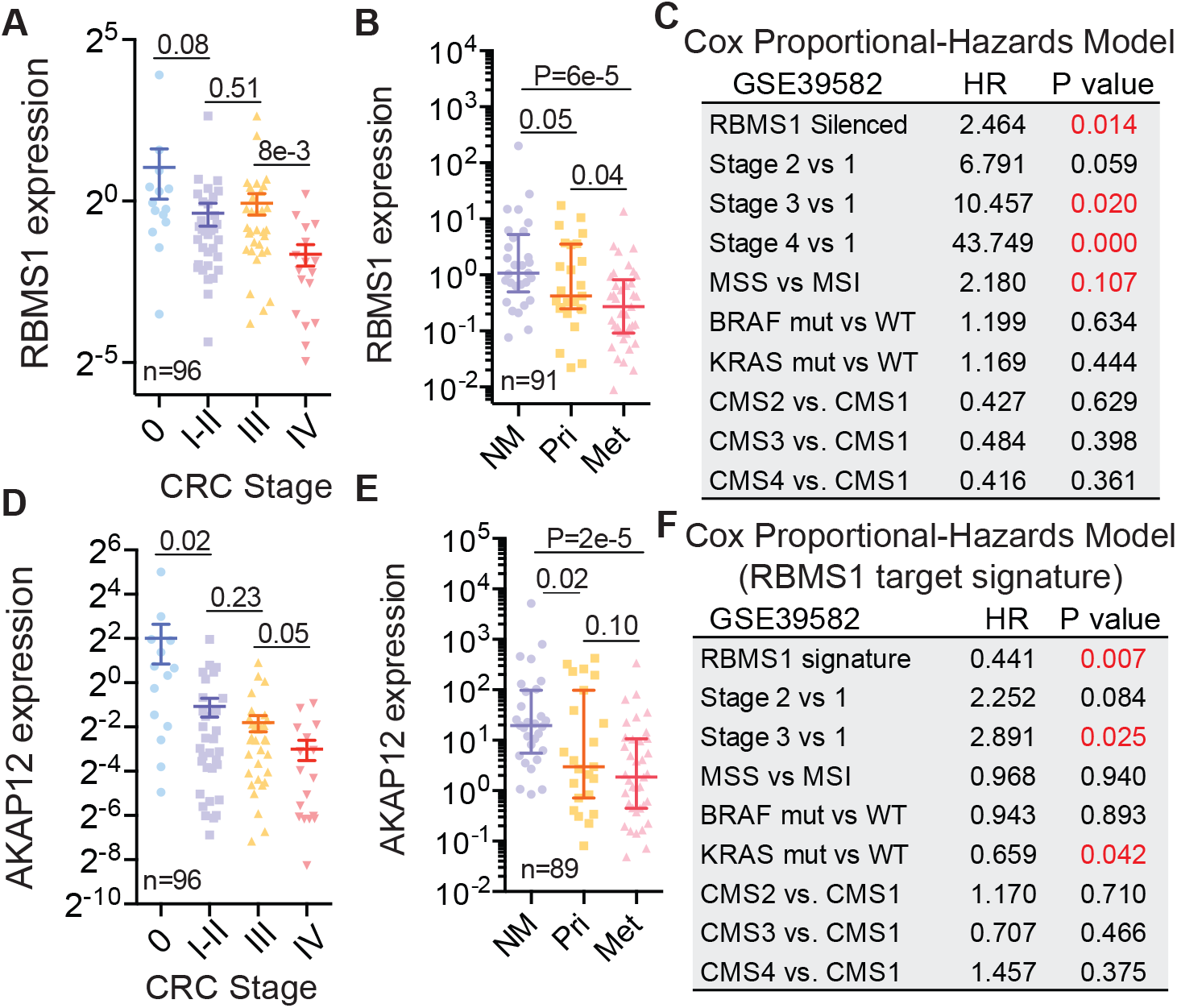
RBMS1 and its regulon are associated with colon cancer metastasis and reduced survival in CRC patients. **(A)** RBMS1 qPCR (relative to HPRT1) in 96 colon cancer samples stratified based on tumor stage. **(B)** RBMS1 qPCR (relative to HPRT1) in 29 normal mucosa, 25 primary colon cancer, and 37 liver metastases. **(C)** Multivariate survival analysis (Cox proportional-hazards model) of colon cancer patients in a publicly available dataset (20) with RBMS1 silencing as one of the factors. Significant *P* values are indicated in red. MSS: microsatellite stable; MSI: microsatellite instable; CMS: consensus molecular subtype. **(D-E)** AKAP12 expression in clinical samples (similar to a and b). **(F)** Survival analysis (Cox proportional-hazards model) for the RBMS1 targets as a gene signature associated with progression.

Our results indicate that AKAP12 acts as a suppressor of metastasis downstream of RBMS1, and therefore is expected to show a similar association with metastatic disease. To test this hypothesis, we performed qPCR measurements in the clinical cohorts mentioned above, and in both cases, we observed a significant reduction in AKAP12 expression as a function of disease progression (Fig. 6D-E). Consistent with RBMS1 acting as a regulator of AKAP12, we observed a highly positive and significant correlation between the expression of these two genes (Supplementary Fig. 6B). Interestingly, our analyses indicate that the identified RBMS1 target genes provide a robust gene signature that is both similar to RBMS1 and is highly informative of clinical outcomes. For this analysis, we defined an RBMS1 target score as an aggregate measure of expression for the entire RBMS1 regulon (average normalized expression of target genes). We then stratified patients based on this score and we observed that lower expression of the RBMS1 regulon is associated with lower relapse-free and overall survival in colon cancer patients (Supplementary Fig. 6C). Consistently, RBMS1 target score remained a significant covariate in a multivariate Cox proportional-hazards model of relapse-free survival (Fig. 6F). The association between RBMS1 regulon expression and CRC metastasis is not limited to this dataset. For example, in line with our initial observations in matched samples (Fig. 1G), comparison of gene expression changes in liver metastases relative to CRC primary tumors in a publicly available dataset (21), revealed a significant reduction in the expression of the RBMS1 regulon (Supplementary Fig. 6D-E). Further analysis of publicly available RNA-seq data using REMBRANDTS, from a set of matched primary and metastatic samples (22), also confirmed (i) RBMS1 silencing in multiple liver metastases and (ii) a concordant reduction in the stability of the putative RBMS1 regulon (Supplementary Fig. 6F). Together, these results establish the clinical relevance of our findings and the importance of RBMS1 silencing both as a prognostic factor and as a suppressor of liver metastasis in patients.

### HDAC1-mediated promoter deacetylation leads to RBMS1 silencing in LS174T cells

Our findings described above establish a previously unknown regulatory pathway driven by the RNA-binding protein RBMS1 that plays a functional role in CRC metastasis to liver. Regulatory pathways can be expanded by uncovering the upstream pathways that influence their activity. We have developed computational and data analytical tools designed to integrate publicly available datasets from a variety of biological sources to identify such upstream regulatory mechanisms (16). Using these tools, we sought to extend the RBMS1 regulatory pathway by exploring the mechanisms for RBMS1 silencing observed in the highly metastatic LS174T colon cancer cells. First, we found RBMS1 expression to be strongly associated with promoter acetylation, but did not identify any specific transcription factors associated with RBMS1 expression (Fig. 7A). Consistent with this observation, analysis of the connectivity map dataset (23), which reports the impact of hundreds of small molecule treatments on gene expression, identified the histone deacetylase inhibitor Trichostatin A (TSA) as an activator of RBMS1 expression and of the RBMS1 regulon (Supplementary Fig. 7A-B). As RBMS1 is endogenously silenced in LS174T cells and expressed in the SW480 line, we tested the relative acetylation levels of the RBMS1 promoter in these two lines by performing H3K27Ac ChIP-qPCR. This data showed relatively low levels of H3K27 acetylation at the RBMS1 promoter in the LS174T line compared to the SW480 line, consistent with the differences in the level of RBMS1 in the two lines (Fig. 7B). TSA simultaneously inhibits multiple HDACs, however, we noted that among this group, HDAC1 is the most significantly upregulated in highly metastatic cells generally, and in LS174T cells compared to SW480 cells in particular (Fig. 7C-D; Supplementary Fig. 7C). Moreover, we observed that RBMS1 and HDAC1 expression are negatively correlated in colon cancer clinical samples (Supplementary Fig. 7D). Furthermore, we observed that silencing HDAC1 increases RBMS1 expression (Fig. 7E; Supplementary Fig. 7E-F). Increased expression of HDAC1, and other class I HDACs, has been reported as a strong predictor of survival in colon cancer patients (24) and modulations in HDAC activity result in widespread gene expression reprogramming, including repression of tumor suppressor genes (25). Our observations here indicate that HDAC1-mediated transcriptional repression may be one possible mechanism for RBMS1 silencing in colon cancer cells. However, other epigenetic mechanisms, such as promoter methylation, may also be used by cancer cells to silence RBMS1 expression *en route* to metastatic progression.

**Figure 7.**
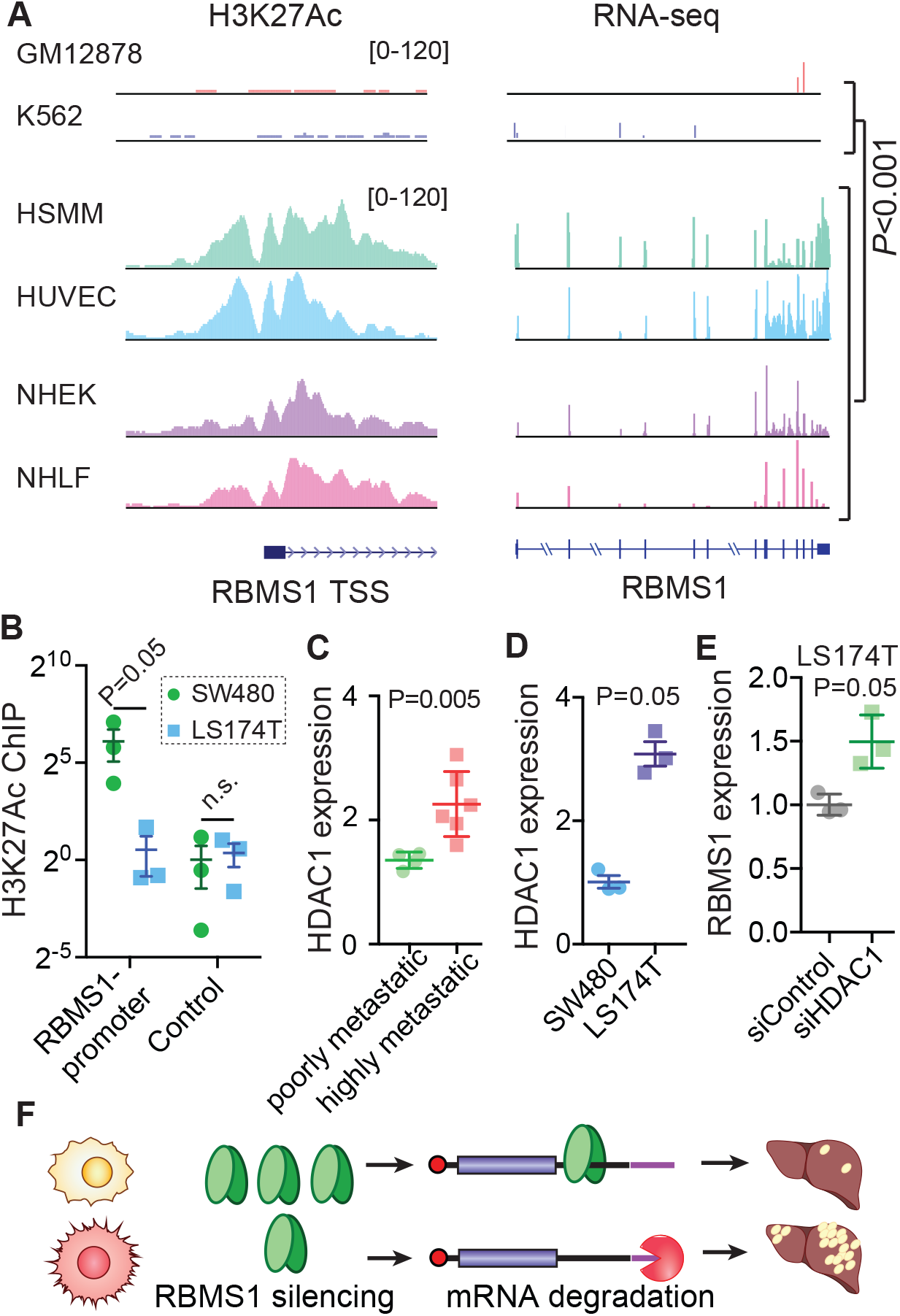
HDAC-mediated promoter deacetylation results in RBMS1 silencing. **(A)** RBMS1 shows dynamic expression and acetylation changes across different cell types. Also shown here is the association between RBMS1 expression and its promoter acetylation (source from ENCODE). **(B)** RBMS1 promoter acetylation levels were measured in SW480 and LS174T cells using H3K27Ac ChIP-qPCR. An unacetylated region ~40kb away from the promoter was used as control (N = 3). **(C)** HDAC1 expression in colon cancer lines stratified based on their metastatic capacity. One-tailed *U-*test was used to compare the two groups. Cell line names are listed in Supplementary Fig. 1A. **(D)** Quantitative PCR to compare HDAC1 levels in the highly metastatic LS174T cells (with silenced RBMS1) relative to poorly metastatic SW480 cells. **(E)** RT-qPCR was used to measure RBMS1 mRNA levels in LA174T cells with RNAi-mediated HDAC1 silencing. **(F)** A schematic model of RBMS1 silencing and its role in suppressing colon cancer metastasis. One-tailed Mann-Whitney *U* tests were used to assess statistical significance for all panels.

## Discussion

Complex human diseases, including cancer, often accompany broad reprogramming of gene expression. In these cases, a comprehensive understanding of the disease state requires not only the identification of the differentially expressed genes, but also understanding the underlying regulatory pathways that explain the dysregulated expression patterns. A number of approaches have been developed by us (11,16) and others (6,26) to tackle this problem. These methods formalize the association between expression and/or activity of master regulators with those of their regulons. Since direct and indirect associations are challenging to disentangle, knowledge of binding preference (*in vitro* or *in vivo*) is also used to better define putative RNA-RBP interactions. Here, we have introduced PRADA to facilitate and formalize the discovery of post-transcriptional regulators that are involved in normal cell physiology and disease. RNA-binding proteins fall into families with highly similar binding preferences (7) and therefore have similar putative regulons. As a result, direct modeling of RBP-target interactions results in unstable predictions (i.e. the model fails to converge on similar outcomes for repeated runs) where the link between a given RBP and its putative regulon is not clear. PRADA solves this problem by integrating the knowledge of changes in the expression of RBPs (used as a proxy for their activities) directly into the analysis. This approach provides a one-step and often stable solution that effectively reveals the RNA-binding proteins whose differential expression is highly informative of changes in gene expression. As RBPs continue to emerge as key regulators of RNA dynamics with crucial roles in human disease, systematic methods like PRADA provide a suitable approach for studying this class of regulators. In this study, we used this approach to discover a previously unknown regulatory network that counteracts colon cancer progression through RBMS1-mediated binding and stabilization of specific metastasis suppressor transcripts.

The association between AKAP12 silencing and colon cancer recurrence and prognosis has been previously described (27). This downregulation is often attributed to AKAP12 promoter methylation and silencing (28). Therefore, identification of the post-transcriptional regulation of AKAP12 mRNA stability by RBMS1 adds another layer of complexity to this regulatory axis in CRC progression. Our analyses of clinical samples presented in Figure 5, however, establish that this association is not limited to AKAP12, and that the entire RBMS1 regulon provides a strong signature for patient survival in CRC. Therefore, it is likely that AKAP12 is not the sole effector of this pathway, and as we showed for SDCBP, other RBMS1 targets may play similar or divergent roles in colon cancer progression.

Given our limited knowledge of the pathways and processes that regulate the RNA life-cycle in the cell, analytical tools that mine quantitative measurements of mRNA dynamics to identify key regulatory interactions can provide an effective avenue for identifying previously unknown molecular mechanisms with critical functions in health and disease. However, these computational strategies must be paired with rigorous experimentation to functionally validate and characterize the putative regulons and their regulators. Using one such framework, we have established a novel functional association between RBMS1 silencing and increased liver metastatic capacity in colon cancer. Our work in the LS174T cell line model indicated that increased activity of histone deacetylase(s), such as HDAC1, could be responsible for RBMS1 silencing. HDAC inhibitors have been used in the clinic to treat metastatic CRC. A number of studies have previously noted increased expression of HDAC1/2 in colon cancer (25) and in cases where RBMS1 silencing also occurs, HDAC inhibitors could be useful as they may also restore RBMS1 expression. While HDAC inhibitors are not specific in their effects, and their effects are not solely mediated through RBMS1, we speculate that in some cases their impact on RBMS1 may lead to better clinical outcomes. Identifying such cases, however, will require a deeper understanding of the mechanisms through which RBMS1 is silenced and the degree to which its expression can be effectively controlled.

## Contributions

H.G. conceived and designed the study. RNA sequencing and analysis was performed by J.Y., J.P.O., and H.G. Cell lines and mouse experiments were carried out by B.C., J.Y., and H.G. B.C performed Immunofluorescent staining. Reporter experiments were conducted by A.N. and B.H. L.F. performed RBMS1 CLIP-seq and western blotting. E.M.W. generated PDX data. PRADA was developed by H.G. and H.A. R.S.W. provided patient tumor samples. Clinical analyses were performed by R.D. and H.G. Manuscript was written by J.Y., L.F., and H.G. H.G. supervised the project.

## Acknowledgments

We acknowledge the UCSF Center for Advanced Technology (CAT) for high throughput sequencing and other genomic analyses. We thank Byron Hann and the Preclinical Therapeutics core as well as the Laboratory Animal Resource Center (LARC) at UCSF. We are also grateful for the genomic data contributed by the TCGA Research Network, including donors and researchers. We acknowledge support from our colleagues at the Helen Diller Family Comprehensive Cancer Center and Rockefeller University. We specifically acknowledge Dr. Sohail Tavazoie for providing access to data from PDX models and for reading an earlier version of this manuscript. We also acknowledge Jonathan Weissman and Luke Gilbert for CRISPRi constructs. We acknowledge Dr. David Erle for providing the backbone construct used for the reporter assay. This work was supported by grants to H.G. (NIH: R00CA194077 and R01CA24098; AACR: 17-20-38-GOOD). J.Y. was supported by an NSF Graduate Research Fellowship, H.A. and L.F. were supported by NIH training grants, F32GM133118 and T32CA108462-15 respectively.

## Supplementary Materials

### Materials and Methods

#### Tissue Culture

SW480 cells were cultured in McCoy’s 5A supplemented with 10% FBS, sodium pyruvate, and L-glutamine. LS174T and 293LTV cells were cultured in DMEM supplemented with 10% FBS, and WiDr cells were grown in EMEM supplemented with 10% FBS.

#### <u>P</u>rioritization of <u>R</u>egulatory Pathways based on <u>A</u>nalysis of RNA <u>D</u>ynamics <u>A</u>lterations (PRADA)

PRADA is a customized variation on lasso regression (least absolute shrinkage and selection operator). Therefore, PRADA can simultaneously perform feature selection and apply a custom penalty function as part of its regularization step. The goal of PRADA is to identify RNA-binding proteins whose differential expression explains changes in the expression of their targets that is observed in the data. We first generated an RBP-RNA interaction matrix based on binding preferences of RNA-binding proteins, as previously reported (7,10). For this, we scanned mRNA sequences for matches to RBP-derived regular expression (as previously described (6)). Given this interaction matrix, our goal is to identify RBPs whose change in expression is predictive of global changes in gene expression of their targets. In other words, we aim to solve the following: Δ*Exp*(*g*) = ∑_*i*_ *α*_*i*_ · *t*_*i,g*_ · Δ*Exp*(*RBP*_*i*_) + *c*_*g*_, where Δ*Exp*(*g*) stands for changes in the expression of gene *g* based on dataset of interest, *t*_*i,g*_ is a binary variable, 1 if *RBP_i_* is predicted to bind *g* and 0 otherwise, Δ*Exp*(*RBP*_*i*_) marks changes in the expression of RBPs themselves as a proxy for their differential activity. The resulting *α*_*i*_ represents the strength of regulatory interactions. To ensure that the most informative RBPs are selected, we introduced a customized penalty term that extends the lasso regression framework: 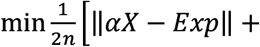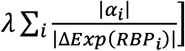. Here, the coefficients are penalized by |Δ*Exp*(*RBP*_*i*_)|^−1^, which is 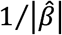 estimate of the linear model used to compare gene expression between the two groups. Other variations of this penalty can also be used, for example standard effect size can account for confidence in 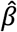 as well 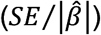. This custom penalty term ensures that RBPs whose activity does not change are not selected by the model. This also stabilizes the resulting model, which would otherwise be a major issue as RBPs that belong to the same family often have very similar binding preferences resulting in correlated features in the interaction matrix. After running this regression analysis, the RBPs with the largest assigned coefficients are prioritized for further study. In this study, we used PRADA to compare poorly and highly metastatic colon cancer lines, which revealed RBMS1 as the strongest candidate. All code and notebooks are available on Github at www.github.com/goodarzilab/PRADA.

#### Stable and transfected cell lines

RBMS1 knockdown cells were using the ViraSafe lentiviral packaging system (Cell Biolabs). shRNA plasmids were obtained from the Sigma-Aldrich TRC library. RBMS1: TRCN0000074936 (sh1), TRCN0000074937 (sh3), and TRCN0000291051 (sh2). AKAP12 and SDCBP were silenced using CRISPR-interference constructs by expressing the following sgRNAs: GGGCGCTCTCGGGACCTCGC and GGCGGCGAGCGGTTCCTTGT. RBMS1 overexpression cells were generated by stably transducing cells with the pLX-304 lentiviral vector containing the RBMS1 open reading frame.

#### Animal studies

All animal studies were performed according to IACUC guidelines. NOD/SCID gamma male mice (The Jackson Laboratory), aged 8 to 10 weeks, were used in all experiments. For splenic (portal circulation) injections, cells were injected directly into the spleen followed by splenectomy (250k for SW480 and LS lines and 100k for WiDr cells). In vivo bioluminescence was measured by retro-orbital injection of luciferin (Perkin Elmer) followed by imaging with an IVIS instrument (Perkin Elmer). For *ex vivo* liver imaging, mice were injected with luciferin prior to liver extraction, and the liver was then imaged and weighed after rinsing with PBS.

#### Quantitative PCR assays

For cell lines, RNA was extracted using the Total RNA Isolation Kit (Norgen Biotek). cDNA was generated using SuperScript III (Invitrogen). SYBR Green Super Mix (QuantaBio) was used to analyze samples on a QuantStudio 6 instrument. Expression of target genes was normalized to HPRT1 expression. Primers sequences are as follows: HPRT1, GACCAGTCAACAGGGGACAT and CCTGACCAAGGAAAGCAAAG; RBMS1, AGGTGCAAAGTCCTTCGTGG and GTGACATGGTGTGCTCCATTG; AKAP12, GAGATGGCTACTAAGTCAGCGG and CAGTGGGTTGTGTTAGCTCTTC; SDCBP, CTGCTCCTATCCCTCACGATG and GGCCACATTTGCACGTATTTCT; eGFP, CCCGACAACCACTACCTGAG and GTCCATGCCGAGAGTGATCC; mCherry, CTGAAGGGCGAGATCAAGCA and TAGTCCTCGTTGTGGGAGGT. For pre-mRNAs, the following primers were used: AKAP12, AGGCCCTGTCGGAGAAA and TCAGAGTCTCTCTGTCCAACTATTA; SDCBP, CTACTTGTTTATGTGTCTGTGGTTAG and CAAGTCTTCGAGAGATGGATAGAG. We achieved 54.8% and 88.6% knockdown for AKAP12 and SDCBP, respectively.

#### Cancer cell proliferation

5×10^4^ cells were seeded into three 6-well wells and subsequently trypsinized and stained with trypan blue to determine cell viability. Viable cells were counted using a hemocytometer at day 3 and day 5. An exponential model was then used to fit a growth rate for each sample (ln(N_t_)~rt where t is measured in days). A linear model was then used to compare growth rates and its associated *p*-values were reported.

#### Cancer cell invasion assays

SW480 shRBMS1 and shCTRL cells were starved for 18h in McCoy’s media with 0.2% FBS. Trans-well invasion assays were then carried out in biological quadruplicates using Corning invasion chambers (354480) as previously described (14).

#### Western blotting

Cell lysates were prepared by lysing cells in ice-cold RIPA buffer (25mM Tris-HCl pH 7.6, 0.15M NaCl, 1% IGEPAL CA-630, 1% sodium deoxycholate, 0.1% SDS) containing 1X protease inhibitors (Thermo Scientific). Lysate was cleared by centrifugation at 20,000 x g for 15 min at 4° C. Samples were boiled in 1X LDS loading buffer (Invitrogen) and 0.1M DTT. Proteins were separated by SDS-PAGE using 4%–12% Bis-Tris NuPAGE gels, transferred to 0.2 mm PVDF (Millipore), blocked using 5% nonfat milk and probed using target-specific antibodies. Bound antibodies were detected using horseradish peroxidase–conjugated secondary antibodies and ECL Western Blotting Substrate (Thermo Scientific), according to the manufacturer’s instructions. Antibodies: beta-tubulin (Proteintech 66240-1-Ig), RBMS1 (Proteintech 11061-2-AP).

#### RBMS1 irCLIP

RBMS1 irCLIP was performed as described in (Zarnegar et al 2016) with some modifications. Briefly, SW480 cells were crosslinked with 400 mJ/cm^2^ 254nm UV. Cells were lysed in lysis buffer (1X PBS, 0.1% SDS, 0.5% sodium deoxycholate, 0.5% IGEPAL CA-630) supplemented with 1X protease inhibitors (Thermo Scientific) and SuperaseIN (Thermo Scientific). Protein-RNA complexes were immunoprecipitated using protein A dynabeads conjugated to anti-RMBS1 (abcam ab150353) for 2 hours at 4° C. Beads were washed sequentially with high stringency buffer, high salt buffer and low salt buffer. Complexes were then nuclease treated on-bead with RNaseI, and then ligated to the irCLIP adaptor using T4 RNA ligase (NEB) overnight at 16° C. RNA-protein complexes were then eluted from beads, resolved on a 4-12% bis-tris NuPAGE gel, transferred to nitrocellulose, then imaged using an Odyssey Fc instrument (LICOR). Regions of interest were excised from the membrane and the RNA was isolated by proteinase K digestion followed by pulldown with oligo d(T) magnetic beads (Thermo Scientific). The resulting RNA was then reverse transcribed using superscript IV (Invitrogen) and a barcoded RT primer, purified using MyOne C1 dynabeads (Invitrogen), and then circularized using CircLigase II (Epicentre). Two rounds of PCR were then performed to first amplify the library using adaptor-specific primers and to add sequences compatible with Illumina sequencing instruments. The libraries were then sequenced on an Illumina HiSeq4000 at UCSF CAT.

#### RNA-seq library preparation

Unless otherwise specified below, RNA sequencing libraries were prepared using RNA that had been rRNA depleted using Ribo-Zero Gold (Illumina) followed by ScriptSeq-v2 (Illumina), and sequenced on an Illumina HiSeq4000 at Center for Advance Technologies (CAT). Matched primary and liver metastases were sequenced using SENSE RNA-seq library preparation kit (Lexogen). AKAP12 knockdown cells were profiled using QuantSeq 3’ mRNA-Seq library prep kit fwd (Lexogen). These libraries were similarly sequenced on an Illumina HiSeq4000 at Center for Advance Technologies (CAT). To quantify and compare gene expression and transcript stability from the SW480 RBMS1 knockdown RNaseq data, first Tophat (v.2.1.1) was used to map the reads to the human transcriptome (build hg38). Cufflinks, cuffmerge and cuffdiff (v.2.2.1) were then used to calculate RPKM and compare gene expression between samples. REMBRANDTS was used to compare changes in RNA stability as previously described. This pipeline was similarly used to study gene expression changes in the CLR PDX models of metastasis. Parental and LvM PDXs CLR4, CLR27, and CLR32 were divided into matched poorly and highly metastatic pairs and analyzed using cuffdiff (for gene expression) and REMBRANDTS (for RNA stability).

To measure RNA stability, SW480 RBMS1 knockdown and control cells were treated with 10 mg/mL a-amanitin (final concentration in the medium). After 9 hours, total RNA was harvested from the cells using the Norgen Cytoplasmic and Nuclear RNA Purification Kit per the manufacturer’s protocol. RNA-seq libraries was then performed and log-fold changes in RNA stability was measured by comparing log(t9/t0) in control and RBMS1 knockdown cells.

#### Immunofluorescence imaging

1.5×10^4^ cells were seeded per well of a 4-chamber slide in 500 uL of culture medium. 48 hours later, the cell culture medium was removed and slides were rinsed 3 times using 500 µL of 1X PBS. Cells were fixed using 400 µL of 4% paraformaldehyde (pH 7.4) for 10 min at room temperature then washed three times with 500 µL of 1X PBS. 400 µL of 0.2% Triton X-100 in 1X PBS was used to permeabilize the cells. After three additional washes, 500 µL of 5% BSA in 1X PBS was added and the cells were incubated at room temperature for 60 min. 1:200 dilution of Rabbit E-cadherin antibody (Cell signaling #3195) in 100µL of 1% BSA was added to the cells and incubated overnight at 4°C. After washing a 1:1000 dilution of fluorescent goat anti-rabbit secondary in 500 µL of 0.1% BSA was added and incubated for 45 min at room temperature protected from light. Samples were washed and air-dried and the mounting medium was added. Slides were then imaged using a Nikon Ti Microscope (Nikon Imaging Center). Images were analyzed using CellProfiler to measure detect cells, measure ECAD signal intensity, and quantify morphological features (v3.1.5).

#### Measurements in clinical samples

For clinical samples, 96 samples across all stages of the disease were acquired from Origene (HCRT104 and HCRT105), 14 normal, 8 stage I, 25 stage II, 32 stage III, and 17 stage IV. HPRT was used as endogenous control, and relative RBMS1 and AKAP12 expression levels were respectively measured across the samples (using the primers listed above). We also extracted RNA from another 100 normal mucosa, primary tumors, and liver metastases. Roughly 90 of these samples yielded sufficient RNA for qPCR, which was carried out as previously described.

#### Clinical association studies

Patients profiled and documented in GSE39582 were first stratified into two groups based on their RBMS1 expression: silenced (bottom 5%) and expressed (otherwise). 5% was selected as the cut-off because it is close to two standard deviations away from the average RBMS1 expression across all samples. A multivariate Cox Proportional Hazard model (R package survival) was then used to evaluate 5-year disease-free survival. Univariate survival analyses, both disease-free and overall, were also carried out with this stratification. For the target regulon, the RBMS1 signature score was calculated as the median expression of target genes in each sample (normalized gene expression data). Disease-free survival was defined as the time from diagnosis to relapse or death due to any cause. We performed Cox Proportional Hazards multivariable modeling using the RBMS1 score as a continuous variable. Variables added to the model included stage, microsatellite status, KRAS and BRAF mutation status, and Consensus Molecular Subtypes.

### Data availability

The data generated as part of this study are accessible through Gene Expression Omnibus (GSEXXXX).

## Supplementary Figures

**Supplementary Figure 1.**
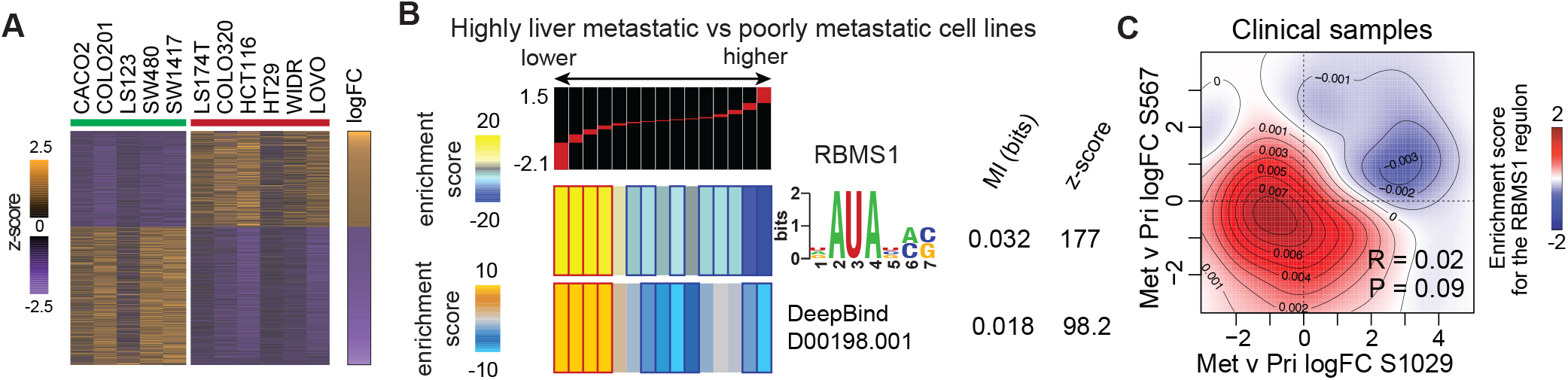
RBMS1 silencing is associated with higher metastatic capacity in colon cancer cell lines, patient-derived xenograft models, and matched primary-metastasis samples from patients. **(A)** Heatmap displaying a gene expression signature associated with CRC cell lines with higher metastatic capacity (GSE59857). Cell lines with lower metastatic capacity indicated by green bar, those with higher metastatic capacity indicated by red bar. **(B)** Similar analysis as (Fig. 1B) but using alternative RBMS1 binding predictions (top: CISBP M143_0.6 (7); bottom: DeepBind model (8)). **(C)** Two-dimensional heatmap showing an enrichment of RBMS1 targets in the group of transcripts that is downregulated in metastases compared to primary tumors from two patients, S1029 and S567. Red indicates RBMS1 regulon enrichment; blue indicates depletion. Also reported are the overall correlations between the two gene expression profiles (Pearson correlation coefficient and the associated *p*-value).

**Supplementary Figure 2.**
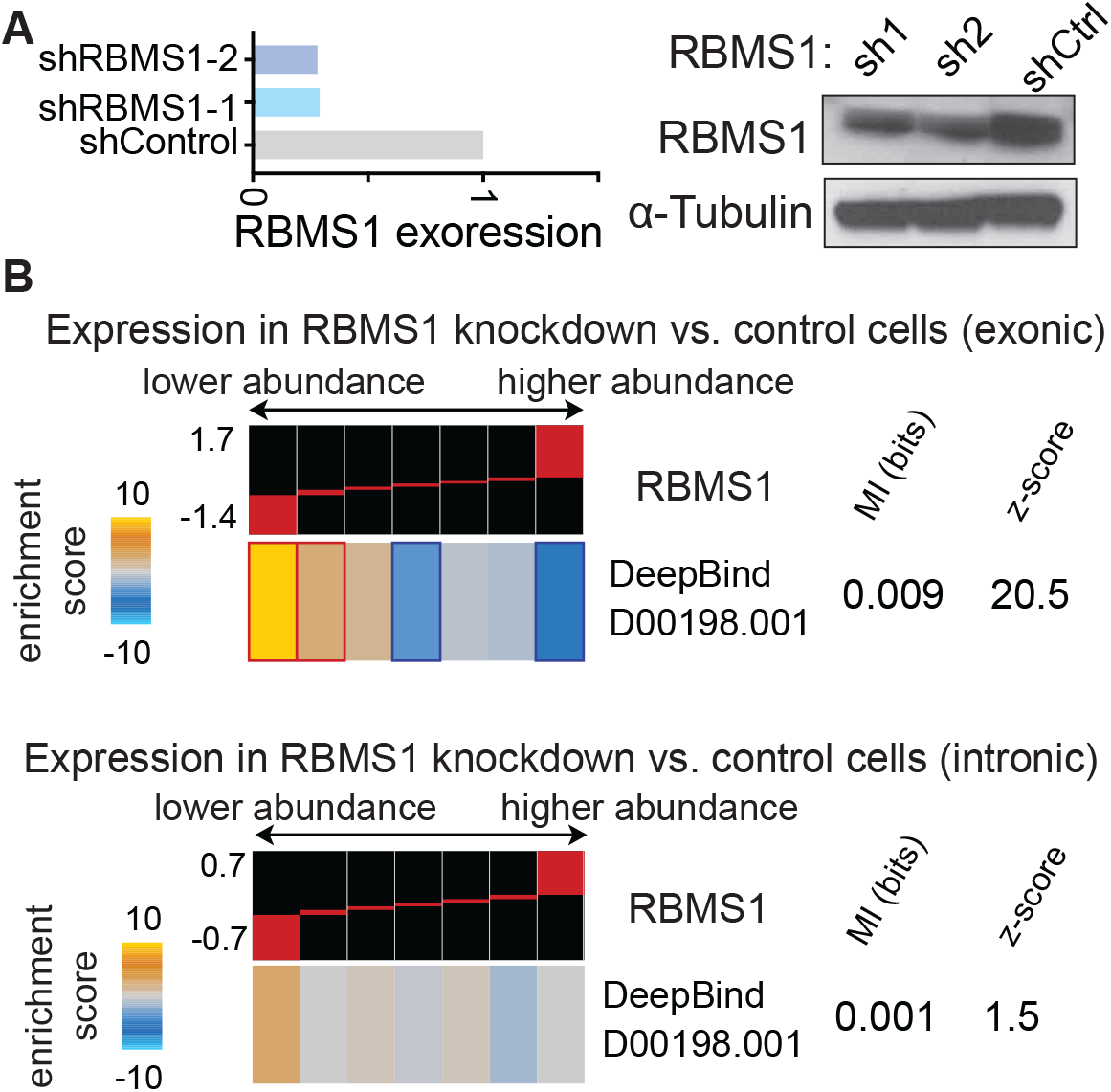
RNAi-mediated RBMS1 silencing and its impact on RNA stability in SW480 cells. **(A)** Western blot confirmation of RBMS1 knockdown using two independent shRNAs in SW480 cells. Tubulin is shown as a normalization control. **(B)** Post-transcriptional changes in RNA stability are reflected in exonic read abundance; however, RNA stability does not impact the abundance of intronic reads (which are correlated with transcriptional changes). Upon RBMS1 silencing, we observed a significant reduction in the expression of the RBMS1 putative regulon as measured by exonic reads (top heat map). However, RBMS1 silencing did not result in the same magnitude of change in RMBS1 regulon expression when measured solely by intronic reads (bottom heat map).

**Supplementary Figure 3.**
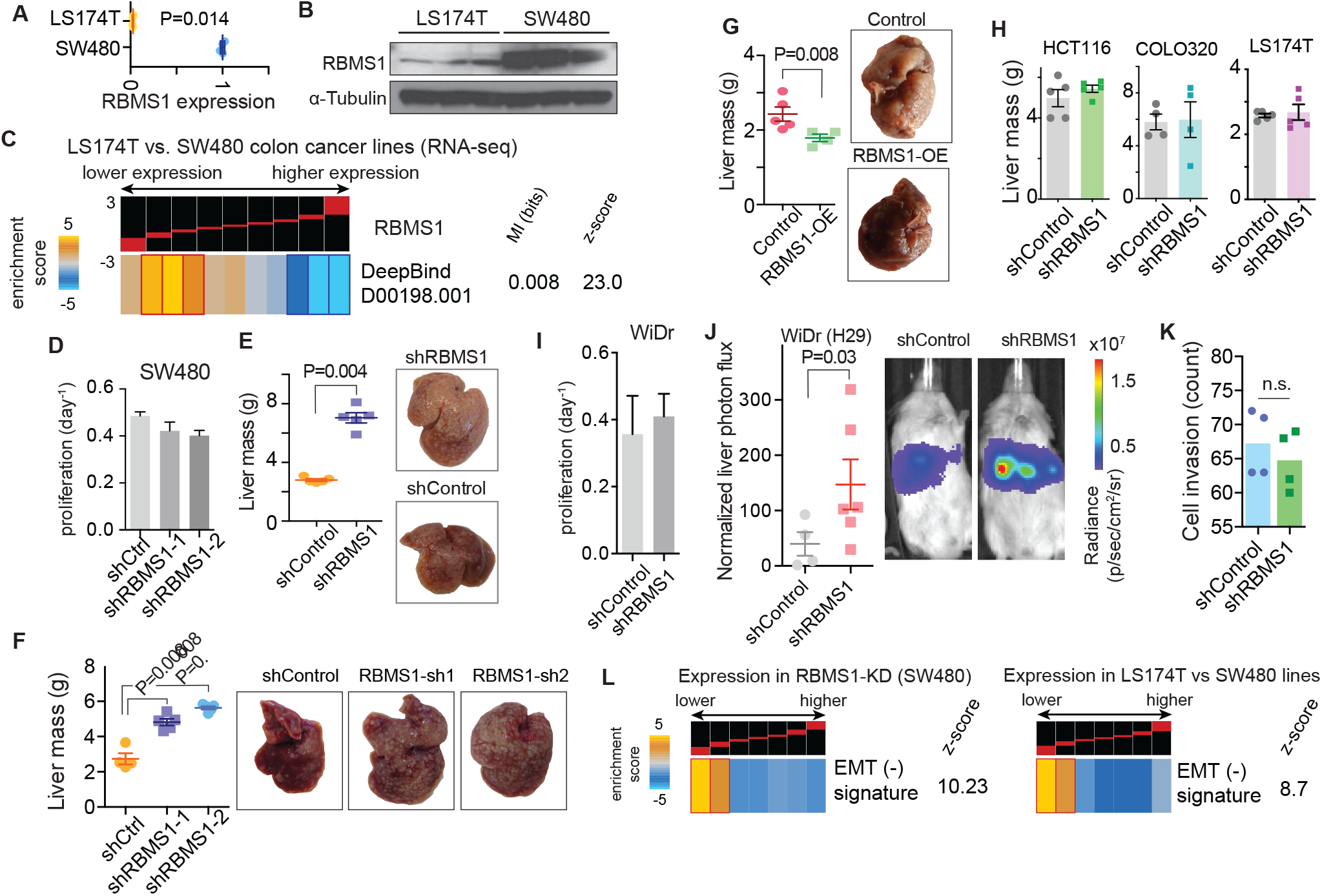
RBMS1 silencing is associated with increased EMT and liver metastasis. **(A-B)** Expression of RBMS1 mRNA (**A**) and protein (**B**), in highly metastatic LS174T and poorly metastatic SW480 cells. One-tailed *U*-test was used to compare qPCR measurements (N = 3). **(C)** Enrichment and depletion patterns of the RBMS1 regulon across gene expression data from RNA sequencing of LS174T and SW480 cells. **(D)***In vitro* cell proliferation rate in RBMS1 knockdown and control cells (two independent hairpins in SW480 cells; N = 3). **(E)** Liver mass as a measure of tumor burden in xenografted mice injected with control or RBMS1 knockdown cells (SW480 background). Also shown are livers collected from representative mice in each cohort. These plots are supplemental to Fig. 3C (N = 5 in each cohort). **(F)** A repeat of the experiment in Fig. 3A (RBMS1 knockdown in SW480 cells) with two independent hairpins targeting RBMS1 (N = 4-5 in each cohort). **(G)** Liver mass as a measure of tumor burden in mice injected with control or RBMS1 over-expression cells (LS174T background). These plots are supplemental to Fig. 3B (N = 5 in each cohort). **(H)** Liver mass measurements (at day 21) in liver colonization assays in HCT116, COLO320, and LS174T lines. RBMS1 is largely silenced in these lines and further knockdown was not expected to elicit any overt change in metastatic colonization. We achieved 60% knockdown in LS174T, 70% in HCT116, and 47% in COLO231. **(I)** *In vitro* cell proliferation rate in RBMS1 knockdown and control cells (WiDr background; N = 3). **(J)** Liver colonization assays (day 49) in RBMS1 knockdown and control cells in an independent line (WiDr/H29; N = 6 and 4, respectively). **(K)** Trans-well invasion assays in RBMS1 knockdown and control cells (SW480) in biological quadruplicates. Statistical significance was assessed using one-sample Mann-Whitney *U* test. **(L)** Heatmaps showing downregulation of EMT(−) signature genes upon RBMS1 silencing, both in LS174T compared to SW480 cells (right), and in SW480 cells with shRNA-mediated RBMS1 knockdown compared to control cells (left). EMT(−) genes are defined as the set of genes, such as E-Cadherin (ECAD), that are downregulated during EMT.

**Supplementary Figure 4.**
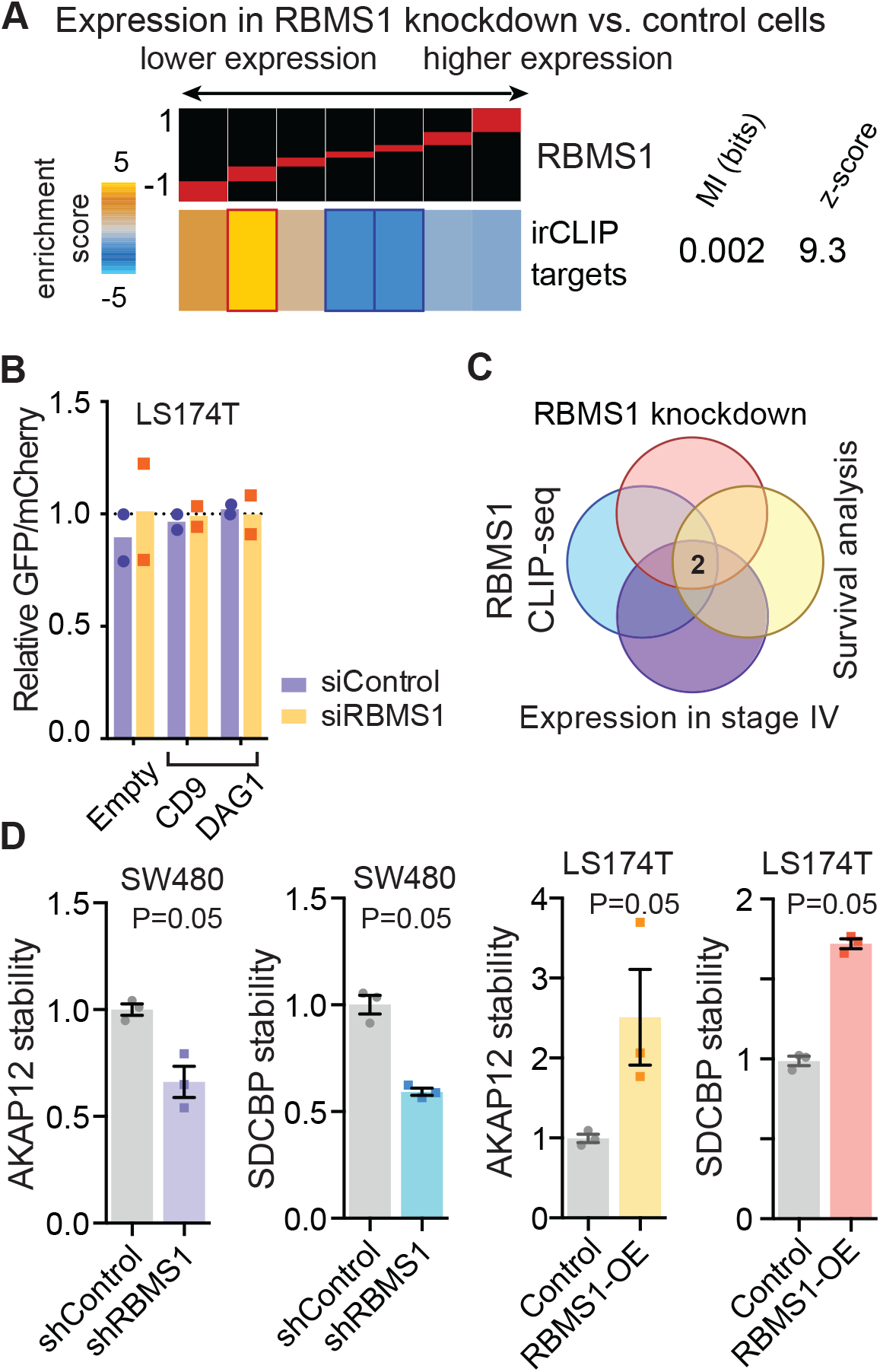
AKAP12 and SDCBP are direct targets of RBMS1. **(A)** Consistent with our results with the putative targets of RBMS1 (DeepBind), CLIP-validated direct targets of RBMS1 in colon cancer cells show a similar decrease in expression and stability in RBMS1 knockdown compared to control SW480 cells. **(B)** The impact of cloning RBMS1 binding sites from specific target genes in the 3’UTR of the GFP/mCherry reporter. Measurements were made in triplicate in both shControl and shRBMS1 expressing cells in the LS174T background. **(C)** An integrated intersection analysis to identify the most robust targets of RBMS1 likely to be involved in metastatic progression. The two genes at the intersection of our four datasets were AKAP12 and SDCBP. **(D)** Changes in the mRNA stability of AKAP12 and SDCBP upon RBMS1 knockdown in SW480 cells and RBMS1 over-expression in LS174T cells.

**Supplementary Figure 5.**
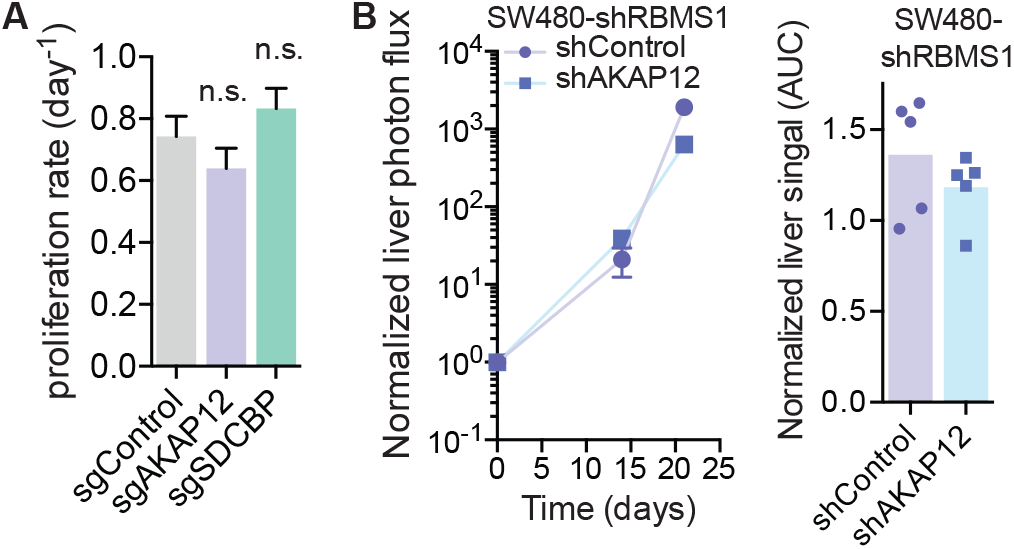
Epistasis experiments confirm the function of AKAP12 downstream of RBMS1. **(A)** Proliferation rates of SW480 cells upon CRISPRi-mediated silencing of AKAP12 and SDCBP. N = 3 in each arm. **(B)** Liver colonization assays were used to compare SW480 cells with knockdown of both RBMS1 and AKAP12 compared to cells with knockdown of only RBMS1. Also shown is the area under the curve (AUC) of log-normalized signal in the liver of the mice over four weeks (N = 5). Further silencing of AKAP12 did not increase liver colonization capacity of cells where RBMS1 was silenced (and therefore AKAP12 levels were endogenously low).

**Supplementary Figure 6.**
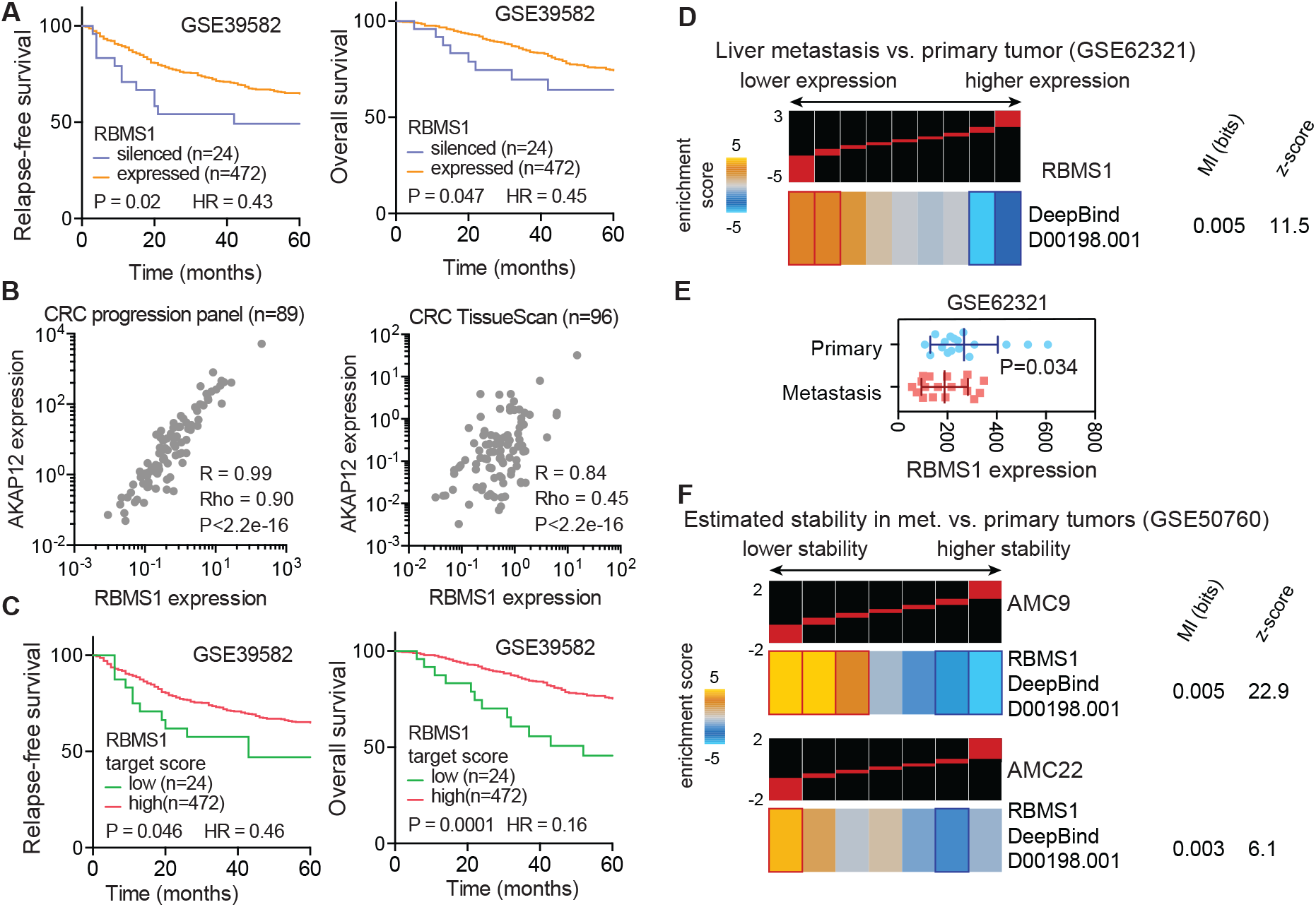
Survival analysis for RBMS1 and its regulon and assessment of its association with metastasis. **(A)** RBMS1 silencing was observed in ~5% of primary CRC tumors (see methods), and these patients showed substantially lower relapse-free and overall survival. Reported are Mantel-Haenszel hazard ratios (HR) and *p*-values from Gehan-Breslow-Wilcoxon tests. **(B)** Regression analysis between AKAP12 and RBMS1 expression levels in two independent clinical cohorts (M = 89 and 96, respectively). Reported are both Pearson and Spearman correlation coefficients (R and Rho) and the associated *p*-value from linear regression analysis. **(C)** Patient primary tumors were scored based on aggregate expression of RBMS1 targets and the resulting values were used to perform survival analyses similar to those in (**A**). As shown here, lower expression of RBMS1 targets were significantly associated with lower relapse-free and overall survival. Also reported are Mantel-Haenszel hazard ratios (HR) and *p*-values from Gehan-Breslow-Wilcoxon tests. **(D-E)** Expression changes in the RBMS1 regulon in liver metastases relative to CRC primary tumors in a publicly available dataset (21). Concomitant with the downregulation of its regulon, RBMS1 itself is also silenced in in this dataset. **(F)** We re-analyzed RNA sequencing of 12 matched CRC primary and liver metastases from publicly available datasets (22). Three patients showed evidence of RBMS1 silencing, and of these, two samples showed a significant reduction in the stability of the RBMS1 regulon (shown as heatmaps).

**Supplementary Figure 7.**
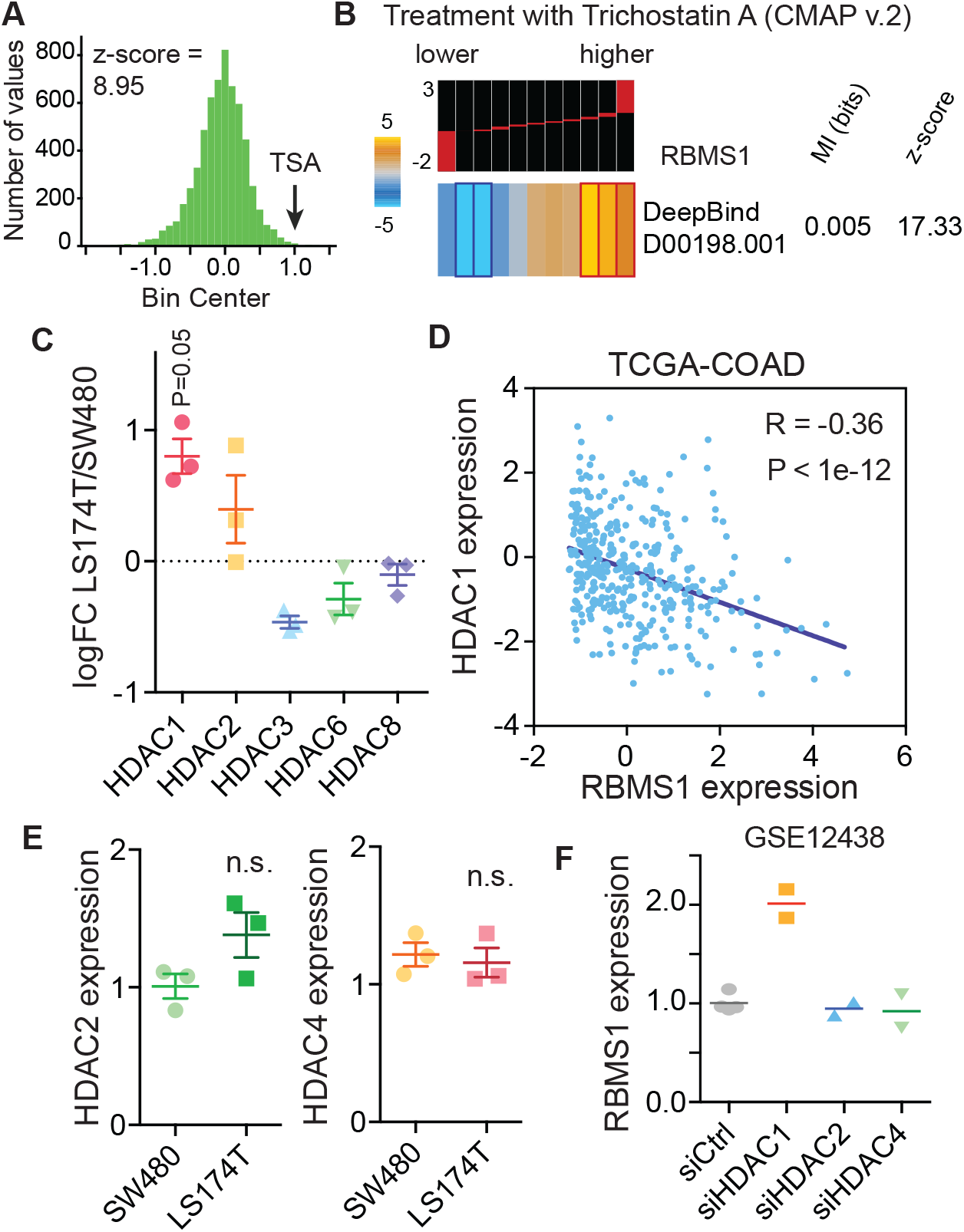
HDAC1-mediated RBMS1 silencing in highly metastatic LS174T cells. **(A)** Distribution of RBMS1 expression across the connectivity map (CMAP) dataset. An arrow indicates a Trichostatin A treated sample and its associated *z*-score. **(B)** Expression of the RBMS1 regulon in HL60 cells treated with Trichostatin A (data from connectivity map), which results in higher RBMS1 expression. **(C)** Comparing the expression of a set of HDACs known to be inhibited by TSA in LS174T cells (where RBMS1 is silenced) relative to SW480 cells. Only HDAC1 shows significantly higher expression in LS174T cells (one-tailed *U-*test). **(D)** Regression analysis comparing the expression of RBMS1 and HDAC1 in TCGA colorectal adenocarcinoma dataset (TCGA-COAD). Shown are the Pearson correlation coefficient and the associated *p*-value. **(E)** Comparison of HDAC2 and HDAC4 expression in SW480 and LS174T cells using qRT-PCR. **(F)** Changes in RBMS1 expression in response to silencing of various HDACs in LNCaP cells (29).

## Notes

**Disclosure of Potential Conflicts of Interest** No potential conflicts of interest are declared by the authors.

